# Mutations in antimicrobial peptides differently affect sleep and plasticity

**DOI:** 10.1101/2025.09.28.679065

**Authors:** Rahul Kumar, Gokul Madhav, Priyanka Balasubramanian, Rhea Lakhiani, Mugdha Joshi, Vibhu Jaggi, Anushna Pal, Imroze Khan, Krishna Melnattur

## Abstract

Molecules implicated in host defense can, independent of their roles as immune mediators, regulate sleep. In mammals, cytokines such as Tumor Necrosis Factor α (TNF α) and Interleukin 1β (IL1β) promote sleep. In *Drosophila*, infection stimulates the systemic release of a battery of anti-microbial peptides (AMPs). These fly immune effectors have been understudied as sleep modulators. We obtained mutants in different AMPs and characterized their functions in sleep and sleep dependent outputs. AMP mutants sleep less with a longer sleep latency, and lowered arousal thresholds. Group C mutant flies, doubly mutant for the anti-fungal peptides Metchinikowin (Mtk) and Drosomycin (Drs), showed the greatest impairments. These Group C mutants also showed an exaggerated sleep rebound. Sleep rebound was unaltered in other mutants. Different sets of AMP mutants exhibited specific disruptions in socialization and rocking induced sleep. These data are a detailed characterization of sleep regulation in AMP mutants. We also evaluated sleep functions. Group C mutants uniquely exhibited normal learning and memory, and lower synapse abundance, despite sleeping the least. Group C mutants are thus able to carry out some sleep functions without sleeping very much. Glial knockdown of Metchnikowin and Drosomycin mimicked the sleep phenotypes of the null mutants, these genes thus act from glia. Sleep and memory defects in AMP mutants was reversible - enhancing sleep of Bomanin mutants pharmacologically or behaviorally improved learning and memory. Together, these data suggest that AMPs are potent sleep modulators and that different classes of AMPs differently affect sleep and sleep dependent outcomes.

## Introduction

Sleep and immunity are intertwined. In humans, sleep loss impairs the immune response – dampening antibody titers induced by vaccines [1] and increasing susceptibility to infections [2]. Conversely, activating the immune system in response to infection or inflammation increases sleep.

Further, in mammals, it is now well established that molecules classically thought to function in immunity can, independent of their role in immunity, regulate sleep. Thus, in classic studies, injection of the cytokines Tumor Necrosis Factor α (TNF α) or Interleukin 1β (IL 1β) led to a dose dependent increase in Non Rapid Eye Movement (NREM) sleep[3]. Conversely, inhibition of TNF α with neutralizing antibodies[4, 5], or loss of function mutations in the TNF α [6] or IL1β receptors [7] decreased NREM sleep. TNF and IL1 are also induced during the acute phase response that follows an infection, concomitant with sleep.

*Drosophila* respond to infection with a robust humoral immune response resulting in the release of a number of potent antimicrobial peptides (AMPs) [8]. These effectors of the fly innate immune response, are released into the hemolymph with acute phase like kinetics from the fat body (functional equivalent of the mammalian liver). They are typically small cationic peptides, regulated by NFΚB transcription factors [9, 10] and the Toll and Immune deficiency cell-surface receptors [11, 12]. In *Drosophila*, a number of AMPs have been classically described - Cecropins, Defensin [13], Attacins [14, 15], Diptericin [16], Drosocin [17], Drosomycin [18], Metchnikowin [19, 20], and Bomanins [21]. A few more AMPs have been recently isolated – Daisho [22], Baramicin [23], and Nemuri [24]. Mutations in the *Drosophila* NFΚB transcription factors *Relish* and *Dorsal related immunity factor (Dif)*, reduce sleep [25, 26]. Surprisingly, studies of the role of the immune effectors, the AMPs, on sleep in flies have been limited, and largely focused on overexpression analysis [24, 27].

A comprehensive, comparative, loss of function analysis of different AMPs and their roles in sleep regulation, the modulation of sleep in different contexts, and sleep function is lacking. Such an approach could be powerful, allowing for the uncovering of generalities as well as particulars (unique roles of specific AMPs) in aspects of sleep regulation and function. Towards this end, we acquired deletions in different AMPs [28]. In immunity, AMPs typically work in combination to combat pathogens. Studies of AMPs in immune function have accordingly evaluated AMPs in combination [28]. For our analysis of AMP functions in sleep, we have followed this convention. AMP deletion mutants are combined into different groups – Group A (mutant for Cecropins and Defensin), Group B (mutant for Attacins, Diptericins, and Drosocin), Group C (mutant for Metchnikowin and Drosomycin), and Bomanins. Importantly, the genetic tractability of the fly model allows us to dissect with precision the roles of different AMPs in aspects of sleep regulation and function. The complexity of mammalian models, in contrast, renders such studies infeasible. Given that connections between molecular players in innate immunity and sleep are common, we expect that our results on fly immune effectors will be broadly applicable.

We find that AMP mutants sleep less, with Group C mutants showing the greatest sleep loss. Different classes of AMP mutants showed differences in sleep rebound (Group C exhibited a hyper rebound) and in sleep plasticity (Group B and Bomanins were impaired in socialization and rocking induced sleep). Strikingly, we found that despite showing the greatest degree of sleep loss, Group C mutants uniquely retained normal learning and memory, and their brains displayed lower synapse abundance. Thus, while many AMPs regulate sleep, sleep functions are uniquely preserved in some AMP mutants (Group C) despite their low sleep. Together, these data are a detailed account of novel roles for these molecules, first identified as effectors of the innate immune response, in sleep and sleep-dependent functions.

## Results

### Mutations in AMPs reduce sleep

To examine the role of antimicrobial peptide (AMP) genes in regulating sleep, we monitored sleep in AMP mutant (Group A, Group B, Bomanin, and Group C) and *iso w*^1118^ control flies. AMP mutant males (Fig 1A and 1B) and females (Fig 1F and 1G) exhibited significantly reduced baseline sleep compared to *iso w*^*1118*^ controls. The extent of sleep suppression differed between the mutants, with Group C mutants showing the biggest effect, and between sexes. Night sleep was significantly reduced in both males and females across all mutants (Fig 1C and 1H). This lowered sleep at night was characterized by shorter sleep bouts (Fig 1D and 1I). In contrast, AMP mutants displayed pronounced sex specific effects on daytime sleep. In males, day sleep time was unaffected in most AMP mutants, apart from Group C mutants, which exhibited shorter day sleep duration (S1A Fig). Sleep consolidation during the day was also not affected in males of most AMP mutants, with the exception of Bomanin mutants, which showed a higher average day sleep bout length (S1B Fig). Females of all AMP mutant lines, however, exhibited lower daytime sleep (S2A Fig). Sleep consolidation during the day was also impaired in females of most AMP mutants, except for Group A, where it was unaffected (S2B Fig).

**Figure 1.**
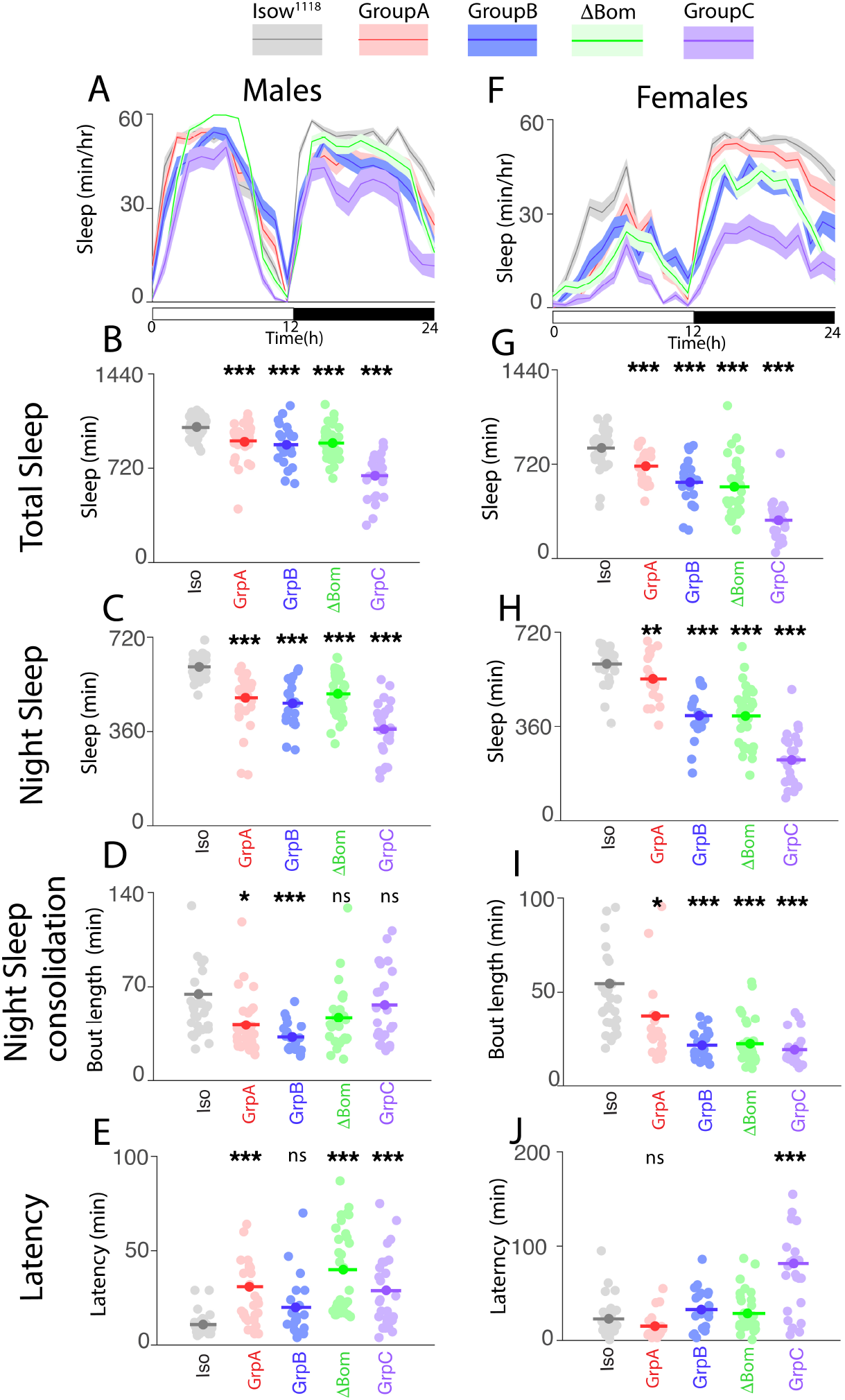
Mutations in AMPs reduce sleep. (A & F) Representative experiment showing sleep (mean ± SEM sleep) per hour of males (A) and females (F) of indicated genotypes. Legend key is for A & F, similar color schemes have been adopted throughout the manuscript. Sleep is shown for 24h, with hours 0-12 as the light period, 12-24 as the dark period. Mean and raw data points are shown for total sleep (B &G), night sleep (C & H), night sleep consolidation (measured by night bout length, D & I), and latency to sleep onset at night (E & J). Sleep characteristics of mutants were compared to *iso w*^1118^ controls. * p<0.05, ** p <0.01, *** p <0.0001. (B & C, G & H) Modified Bonferroni correction following one-way ANOVA for genotype. D & E, I & J, Dunn’s multiple comparisons following Kruskal-Wallis ANOVA (n=20-32 flies/genotype). Iso = *iso w*^1118^, GrpA = Group A, GrpB = Group B, ΔBom = Bomanins, GrpC = Group C.

The probability of falling asleep (pDoze) and waking up (pWake) [29] during the day and night also revealed interesting sex-specific patterns. In males, p[Wake] and p[Doze] largely paralleled the sleep data. Males of all AMP mutant groups showed significantly reduced p[Doze] and increased p[Wake] at night (S1D and S1F Fig), consistent with the broad reductions in nighttime sleep in AMP mutant males. p[Doze] was reduced during the day in Group C mutant males, and unaffected in other AMP mutants during the day (S1C Fig); p[Wake] was unaltered in all of the AMP mutants during the day (S1E Fig). Of note, in males, the sleep of Group C mutants was uniquely reduced during the day. In contrast, the sleep of females was characterized by increased p[Wake] both during the day and night (S2E and S2F Fig). p[Doze] was largely unaffected in females except for Group B and Group C mutants, where it was reduced at night (S2D Fig), and the Bomanin mutants, which exhibited an anomalous increase in p[Doze] during the day (S2C Fig). The broad reductions in sleep time in AMP mutants were accompanied by an increase in time to initiate sleep (sleep latency), particularly in males (Fig 1E). Sleep latency trended higher in Group B and Bomanin mutant females but was only significantly altered in Group C mutant females, where it was dramatically higher (Fig 1J). Importantly, no consistent changes were seen in waking activity. Waking activity was unchanged in AMP mutant females (S2G Fig). Males from Group A and Group B mutants showed slightly but significantly higher waking activity, waking activity was unchanged in Bomanin and Group C mutant males (S1G Fig).

Across taxa, alterations in arousal thresholds are a defining characteristic of sleep [30, 31]. Changes in arousal thresholds are generally regarded as a marker of sleep depth. *Drosophila* also exhibit distinct sleep stages typified by changes in arousal thresholds [32, 33]. To determine the extent to which sleep depth was altered in AMP mutants, we evaluated arousal thresholds using the sensitive *Drosophila* Arousal Tracking (DART) system [34]. In females, arousal thresholds were generally lower in AMP mutants, except for Group A mutants, which exhibited a slight but significant increase in arousability during the day (S3 Fig). In males, AMP mutants exhibited lower arousal thresholds during the day except for Group A mutant males, where arousability during the day was unchanged (S3 Fig). Group C mutant males uniquely exhibited a lower arousal threshold at night. Arousability at night was unchanged in Bomanin mutant males, and slightly but significantly increased in Group B and Group A mutant males (S3 Fig). One possibility to explain these results is that while sleep time is reduced in these mutants, the sleep they do obtain is deeper. Another possibility is that sleep and arousability are (partially) uncoupled in these mutants [35]. Further experiments should help distinguish between these possibilities.

Across all sleep traits and between sexes, the strongest and most consistent effects were seen in Group C mutants. All the AMP mutant lines have been outcrossed into the control *iso w*^1118^ background [28]. Thus, genetic background is not a significant contributing factor in the phenotypes we observe. To help reinforce the specificity of the AMP deletion mutant lines, we employed a whole-body knockdown approach using RNAi driven by a ubiquitous *Da-GAL4* driver. Many of the mutant lines we analyzed carry lesions in numerous genes. The *Bom*^Δ 55c^ allele is a deletion that takes out 10 Bomanin genes [21]. Group B combines deletions in 4 Attacins, 2 Diptericins, and Drosocin. Group A is a composite of deletions in 3 cecropins and Defensin. Group C, in contrast, contains mutations in just two genes – *Metchnikowin (Mtk)* and *Drosomycin (Drs)*, making it tractable to analyze by RNAi-mediated knockdown. The *Da-GAL4 > UAS-Drs RNAi; UAS Mtk RNAi, UAS-Dcr2* flies exhibited substantially reduced sleep during both day and night relative to *Da-GAL4 / +* and *> UAS-Drs RNAi; Mtk RNAi, UAS-Dcr2 / +* parental controls (S4A – S4C Fig). Sleep consolidation was also impaired, and sleep latency increased, without affecting waking activity (S4D – S4G Fig). p[Wake] was increased and p[Doze] decreased, consistent with the changes in sleep (S4H – S4K Fig). RNAi-mediated knockdown of *Drs* and *Mtk* thus closely resembled the sleep deficits observed in Group C mutants (S4 Fig).

### Rebound sleep is enhanced in Group C mutants

AMP mutants sleep less (Fig 1). We hypothesized that the low sleep was a result of these mutants not being able to generate the sleep they need. A natural prediction of this hypothesis is that the mutants would be impaired in the sleep rebound following sleep deprivation. Therefore, to test our hypothesis, we examined the sleep rebound following sleep deprivation in AMP mutants and *iso w*^1118^ controls (Fig 2). We sleep deprived flies overnight using the Sleep Nullifying Apparatus (SNAP) [36, 37] and evaluated rebound over 48h following the period of sleep deprivation. In this paradigm, *iso w*^1118^ controls were effectively deprived of >98% of their sleep (Fig 2A) and recovered ∼50% of the sleep lost over 48h (Fig 2A and 2B). Group A, Group B, and Bomanin mutants did not exhibit any changes in sleep rebound over 48h compared to *iso w*^1118^ controls (Fig 2C-2E). Group C mutants, which slept the least, surprisingly exhibited an exaggerated rebound of ∼91% (Fig 2F and S5A). Thus, contrary to our prediction, none of the mutants exhibited an impaired sleep rebound.

**Figure 2.**
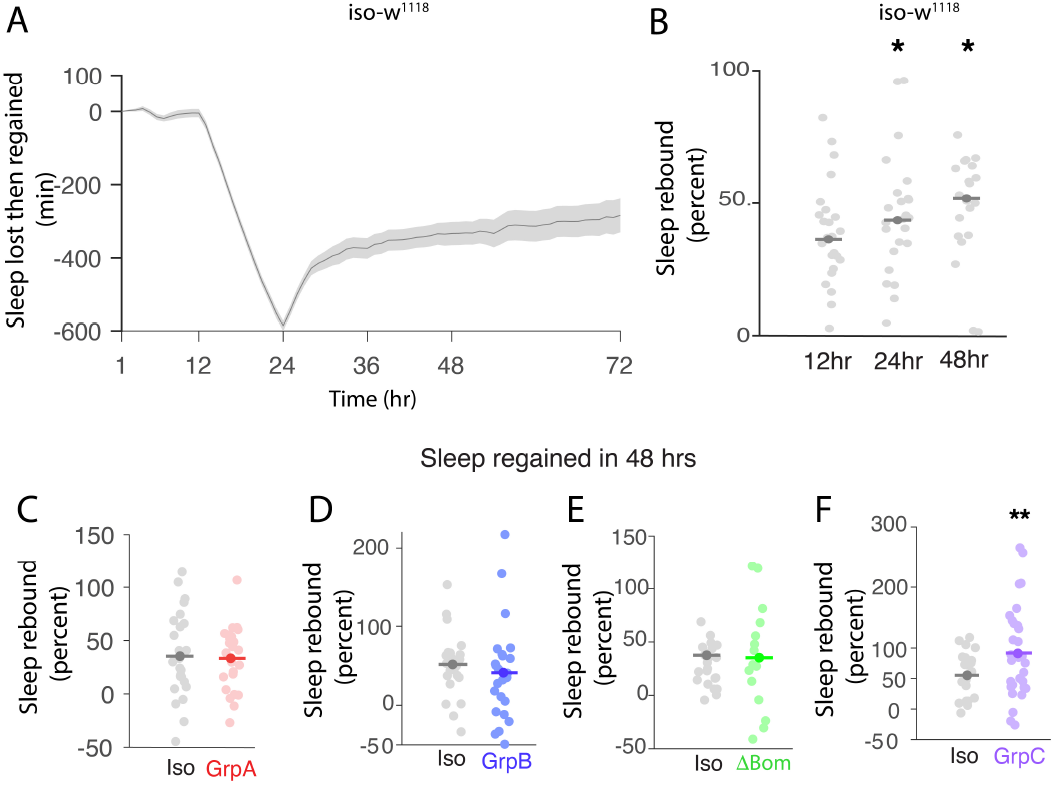
Group C mutants exhibit an exaggerated sleep rebound. (A) Plot of mean ± SEM sleep lost during overnight sleep deprivation and gained during 48h of recovery for *iso w*^1118^ controls. Repeated measures ANOVA, *** p<0.0001. (B) Mean and raw data points are shown for percent sleep regained in 12, 24, and 48 hours of recovery for *iso w*^1118^ controls. (C-F) Recovery sleep over 48 hours of indicated mutants and *iso w*^1118^ controls. Mean and raw data points are shown. (B) One Way ANOVA, followed by Bonferroni correction, (C-F) Student’s T-test (n=24-32 flies/genotype). * p<0.05, ** p <0.01, *** p <0.0001. Iso = *iso w*^1118^, GrpA = Group A, GrpB = Group B, ΔBom = Bomanins, GrpC = Group C.

A close examination of sleep rebound dynamics, however, revealed an interesting pattern. Group C and Group B mutants were impaired in the extent of sleep rebound after the first 12 hours of recovery, both exhibiting approximately half the recovery sleep observed in *iso w*^1118^ control flies (S5B – S5D Fig). Group B mutants eventually caught up to match the *iso w*^1118^ control recovery sleep level as measured after 48h of recovery (Fig 2D). Group C mutants continued to recover lost sleep over the two nights following the period of sleep deprivation to ultimately exhibit an exaggerated sleep rebound over a 48-hour recovery period (S5A and S5B Fig). Importantly, knockdown of *Mtk* and *Drs* with RNAi pheno-copied the results above with the Group C mutant.

*Da-Gal4 > UAS-Drs RNAi; UAS Mtk RNAi, UAS-Dcr2* flies also exhibited a hyper sleep rebound relative to *Da-Gal4 / +* and *> UAS-Drs RNAi; Mtk RNAi, UAS-Dcr2 / +* parental control flies (S5E and S5F Fig).

Sleep is regulated by the coordinated action of two processes – Process S, a homeostatic process, and Process C, a circadian process [38]. Having evaluated sleep homeostasis, we also examined circadian parameters in AMP mutants. *iso w*^1118^ controls exhibited strong behavioral rhythmicity in constant darkness with a period length of 24.0 h (Fig 3A and 3F). Group C mutants exhibited a minor but significant shortening of period length (23.75 h) (Fig 3E and 3J), period length was unchanged in the other AMP mutants (Fig 3B – 3D and 3G – 3I). Circadian behavior in flies is also reflected in anticipatory increases in locomotory behavior before light transitions. We quantified anticipatory behavior using the method of Harrisingh [39] (S6 Fig). Group B (S6D and S6H Fig) and Group C mutants (S6F and S6J Fig) displayed reduced morning and evening anticipation; Bomanin mutants were unchanged in morning anticipation but increased evening anticipatory activity (S6 Fig). These data provide evidence that AMPs are required for aspects of circadian behavior and suggest new avenues for more mechanistic studies. However, the reductions in circadian period we observed in particular are mild, especially in contrast to the large changes in sleep (Fig 1).

**Figure 3.**
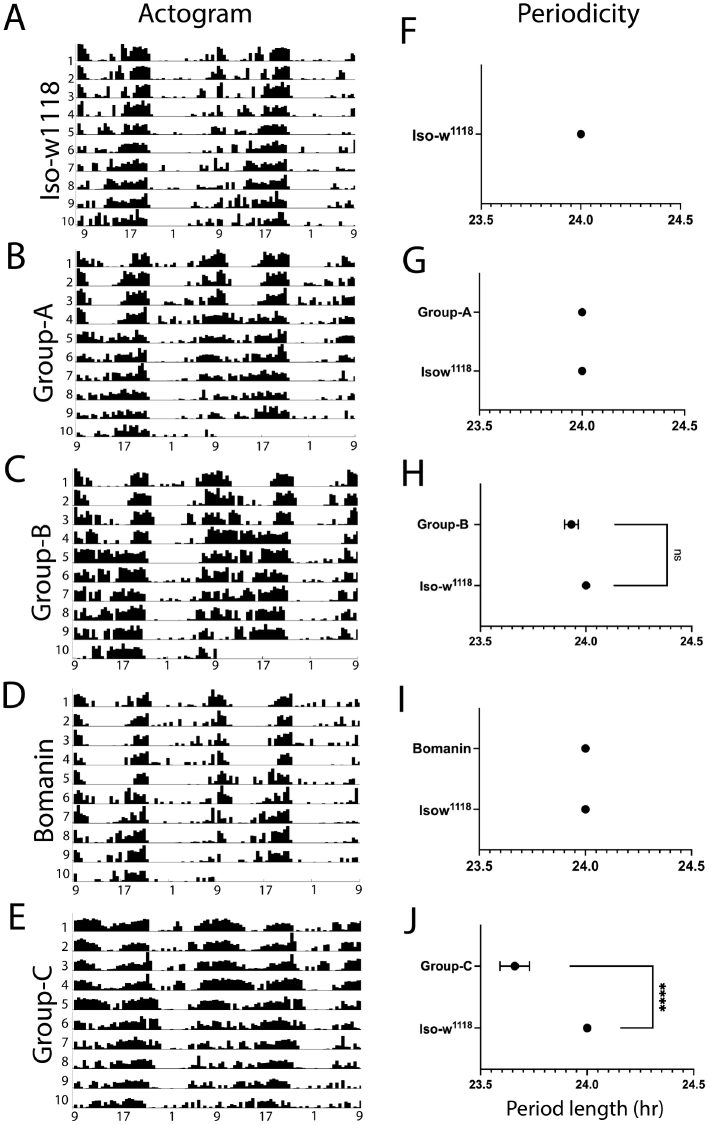
Group C mutants exhibit a shorter circadian period. (A-E) Representative double-plotted actogram showing activity and rest for 10 days of *iso w*^1118^ controls, Group A, Group B, Bomanin, and Group C mutants. Days 1-4 were 12L-12D, and days 5-10 were 12D-12D. (F-J) The mean period length of AMP mutants was compared to *iso w*^1118^ controls. (E & F) Mann-Whitney, two-tailed test (n=26-32 flies/genotype), **** p <0.00001.

### Group B and Bomanin mutants are impaired in socialization and rocking induced sleep

AMP mutants exhibit changes in sleep and sleep homeostasis (Fig 1 and 2). Across many diverse animals, sleep is plastic and modifiable in response to environmental conditions and ecological niches [30, 40]. In flies, sleep is known to be modifiable by socialization [41-43], rocking [44, 45], starvation [46, 47], sexual arousal [48, 49], flight disruption [50], and learning [51, 52]. Understanding how AMP mutants contribute to how sleep is altered in these different circumstances would be expected to provide a much deeper understanding of the roles of AMPs in sleep regulation than could be obtained from a study of baseline sleep alone.

We started by evaluating the changes in sleep following socialization (Fig 4A). Social enrichment for five days resulted in a modest but significant increase in daytime sleep in control *iso w*^1118^ flies compared to their non-enriched siblings kept in isolation post-eclosion (S7A Fig). We calculated the change in daytime sleep in AMP mutants and normalized it to the change seen in *iso w*^1118^ controls (S7B and S7C Fig). Group B and Bomanin mutants exhibited impairments in socialization induced sleep in both males and females (Fig 4B and 4C). In contrast, Group A mutant females, but not males, showed impairments (Fig 4B and 4C). Thus, the effect of AMP loss of function on socialization induced sleep is influenced by both genotype and sex.

**Figure 4.**
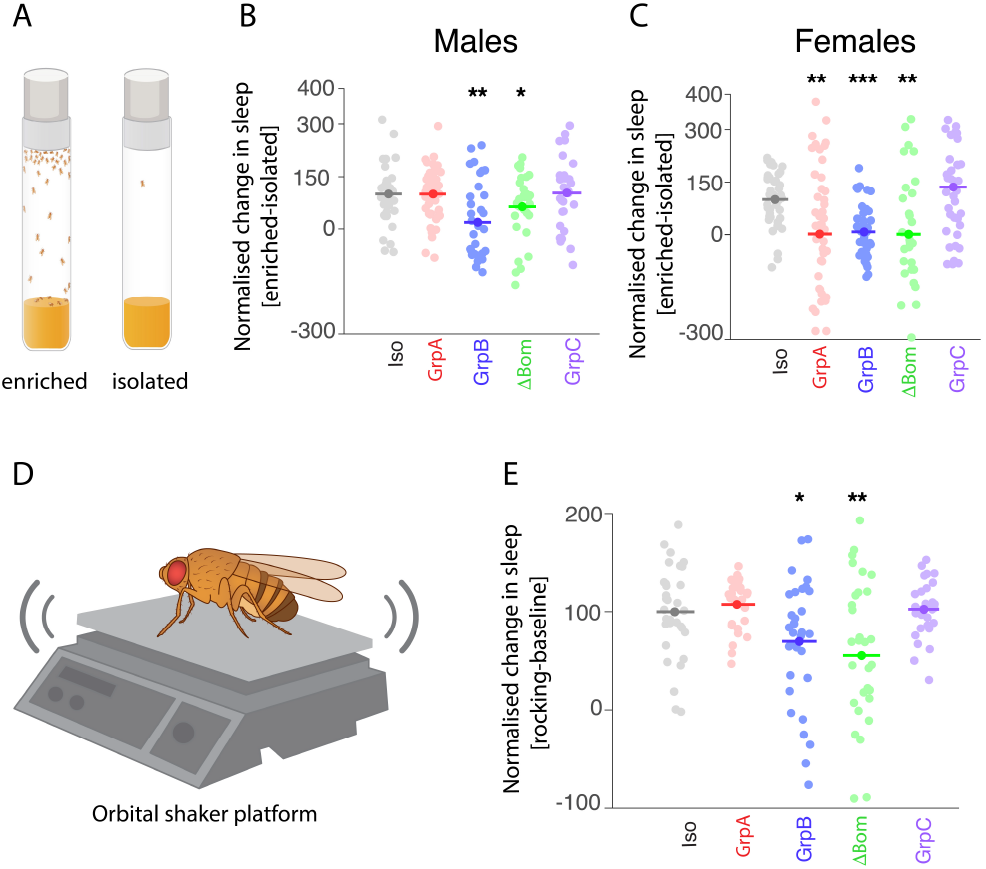
Group B and Bomanin mutants are impaired in socialization and rocking-induced sleep. (A) Schematic of the social enrichment assay. (B & C) Normalized change in sleep upon social enrichment in *iso w*^1118^ controls and AMP mutant males (B) and females (C). (D) Schematic of the rocking assay. (E) Normalized change in sleep upon rocking in *iso w*^1118^ controls and AMP mutant females. Mean and raw data points are shown. Sleep changes in mutants were compared to *iso w*^1118^ controls. * p<0.05, ** p <0.01, *** p <0.0001. (B, C & E) Modified Bonferroni correction following One-way ANOVA for genotype (n=22-32 flies/genotype). Iso = *iso w*^1118^, GrpA = Group A, GrpB = Group B, ΔBom = Bomanins, GrpC = Group C.

Mechanical stimulation by gentle rocking during the day has been shown to increase daytime sleep in flies [44, 45]. *iso w*^1118^ flies displayed a robust increase in daytime sleep (S7D Fig) when subjected to gentle rocking at 90rpm on an orbital shaker (schematic in Fig 4D). We calculated the change in daytime sleep in AMP mutants and normalized it to the change seen in *iso w*^1118^ controls (S7E and S7F Fig). As with socialization induced sleep, Group-B and Bomanin mutants also displayed impairments in rocking-induced sleep (Fig 4E); other AMP mutants appeared unaffected.

### Sex specific changes in starvation induced sleep suppression in AMP mutants

Starvation suppresses sleep in animals from flies to humans [46, 47, 53]. *Drosophila*, when placed on starvation media for 24h (Fig 5A), show robust sleep suppression. Consistent with previous results, *iso w*^1118^ flies also robustly suppressed their sleep when starved for 24h (S8A Fig). We calculated the extent of starvation induced sleep suppression as the percentage of total sleep lost on the day of starvation relative to the baseline day. *iso w*^1118^ flies lost ∼40-50% of sleep upon starvation (S8C Fig). AMP mutants displayed marked sex specific alterations in the extent of starvation induced sleep suppression. Group A mutant males, but not females, were impaired in starvation induced sleep suppression (S8B and S8C Fig), i.e., they lost less sleep on starvation (Fig 5B and 5C). Group C mutant females, but not males, were impaired in starvation induced sleep changes. Bomanin mutant females, but not males, showed exaggerated response to starvation i.e., they lost more sleep (Fig 5B and 5C). Group C mutants are the shortest sleeping AMP mutants. Sleep of Bomanin mutants under starvation though was substantially lower than that of Group C mutants. The impairments in starvation induced sleep suppression in Group C mutants are thus unlikely to be explained by a ‘floor’ effect of low baseline sleep in Group C mutants.

**Figure 5.**
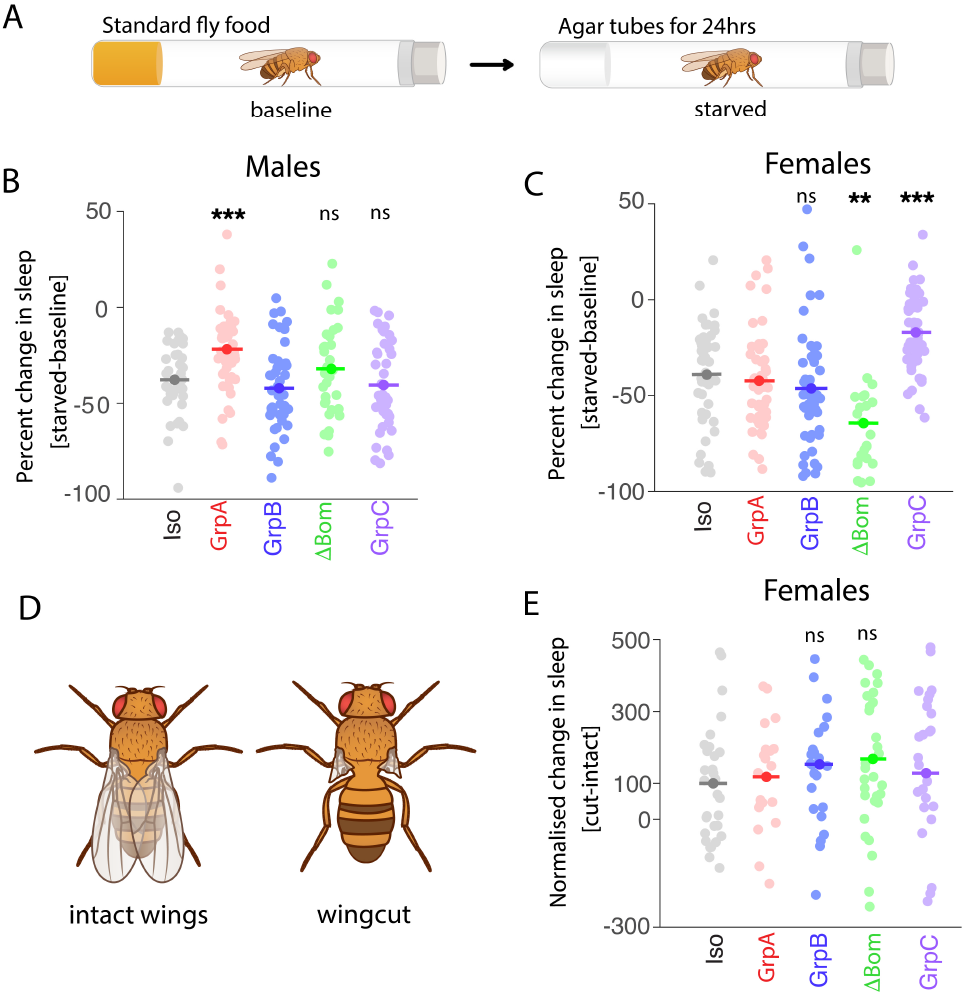
AMP mutations exhibit sex-specific changes in starvation induced sleep suppression. (A) Schematic of the starvation assay and protocol. (B & C) Normalized change in sleep upon starvation in *iso w*^1118^ controls and AMP mutant males (B) and females (C). Mean and raw data points are shown. (D) Schematic showing a fly with intact wings and one with wings cut. (E) Normalized change in sleep upon wing-cut in *iso w*^1118^ controls and AMP mutant females. Mean and raw data points are shown. (B, C & E) Modified Bonferroni correction following Oneway ANOVA for genotype (n=22-32 flies/genotype). Iso = *iso w*^1118^, GrpA = Group A, GrpB = Group B, ΔBom= Bomanins, GrpC = Group C. ** p <0.01, *** p <0.0001

Group B mutant males and females showed a normal response to starvation when considering total sleep over 24 hours. However, analysis of daytime and nighttime sleep changes revealed perturbations. Group B mutant males and females exhibited an exaggerated sleep suppression during the day (S8D and S8E Fig). Group B mutant males, but not females, also showed an impaired sleep suppression at night (S8D Fig). These day vs night specific changes in sleep following starvation would have been missed if sleep were only evaluated over 24 hours.

*Drosophila* suppress sleep when put on starvation media but only recover a small fraction of the lost sleep when put back on regular food. Having evaluated the extent of sleep suppression in AMP mutants, we next evaluated the extent of the lost sleep recovered in AMP mutants and *iso w*^1118^ controls. We quantified the sleep recovered post starvation as the change in sleep on the recovery day compared to the baseline day. *iso w*^1118^ flies recovered about 10% of the sleep lost during starvation (S8F – S8H Fig). Group A and Group C mutants displayed increased recovery sleep post-starvation compared to controls; other AMP mutants were unchanged (S8H – S8J Fig).

In flies, flight disruption is known to increase sleep as a form of sleep-plasticity [50]. We disrupted flight by cutting the wings (schematic in Fig 5D) and evaluated sleep two days after the injury as per established protocols [50]. *iso w*^1118^ flies showed a modest but significant increase in sleep (S8K Fig). We quantified the change in total sleep upon wing cut in AMP mutants and normalized it to the change seen in *iso w*^1118^ controls. The change in flight-disruption induced sleep trended higher in Group B and Bomanin mutants but did not approach statistical significance; other AMP mutants were unaffected (Fig 5E and S8L Fig).

### Lower synapse abundance in Group C mutant brains

These results are a comprehensive characterization of changes in sleep regulation in AMP mutants. We next wanted to evaluate whether AMP mutants exhibit changes in sleep functions. An influential theory of sleep function is the synaptic homeostasis hypothesis (SHY) [54]. As per SHY, waking increases and potentiates synapses, changes which are reversed by sleep. In *Drosophila*, immunostaining with the anti-Bruchpilot (anti-Brp) antibody has been used to quantify changes in synapse abundance [55, 56].

Consistent with previous results, and consistent with the predictions of SHY, we found that brains of sleep deprived *iso w*^1118^ controls exhibited greater Brp staining (S9 Fig). We then proceeded to examine Brp levels in brains from AMP mutants and *iso w*^1118^ controls, reasoning that the increased wakefulness seen in AMP mutants, should reflect in changes in Brp levels as per SHY. Surprisingly, Group C mutant brains showed lower Brp staining despite their exhibiting the greatest sleep loss (Fig 6G and 6H). Brains of other AMP mutants did not show significant changes in Brp intensity (Fig 6A. – 6F).

**Figure 6.**
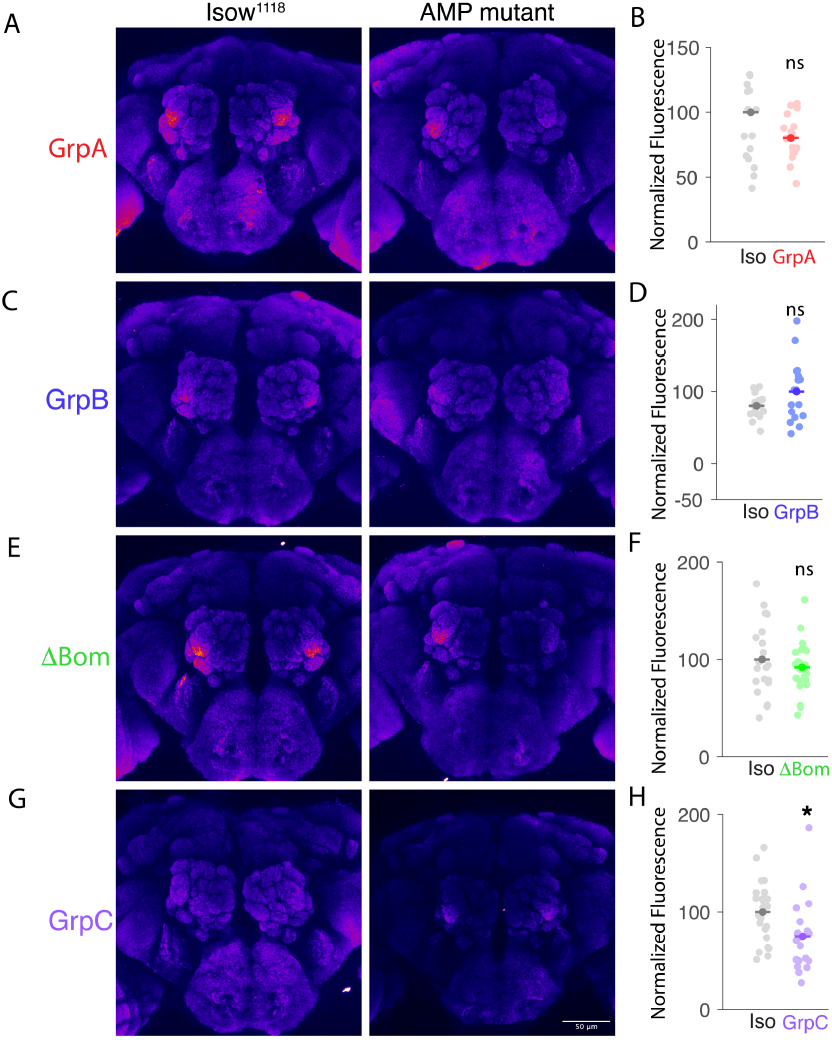
Group C mutant brains exhibit lower synapse abundance. (A, C, E, G) Brains from Group A (A), Group B (C), Bomanins (E), and Group C (G) flies and paired *iso w*^1118^ controls stained with mAB nc82. In each panel, the left image is of an *iso w*^1118^ control brain, and the right image is of an AMP mutant brain. (B, D, F, H) Quantification of BRP expression in Group A (B), Group B (D), Bomanin (F), and Group C (H) brains compared to *iso w*^1118^ controls. Mean and raw data points of normalized fluorescence are shown. (A, C, E, G,) z-projections from whole mount immunostaining processed using the Fire LUT in FiJi. Scale bar in G = 50μm. All images are taken at the same resolution. * p<0.05, ** p <0.01, *** p <0.0001. B, D, F & H-Unpaired t test, two-tailed (n= 17-23 brains/genotype). Iso = *iso w*^1118^, GrpA = Group A, GrpB = Group B, ΔBom = Bomanins, GrpC = Group C.

### Group C mutants uniquely retain normal learning and memory

Sleep loss impairs learning and memory in organisms from nematodes to humans [37, 57-61]. To investigate whether sleep loss in AMP mutants leads to cognitive defects, we assessed short-term learning and memory in flies using the well-established aversive gustatory conditioning assay (schematic shown in Fig 7A) [62].

**Figure 7.**
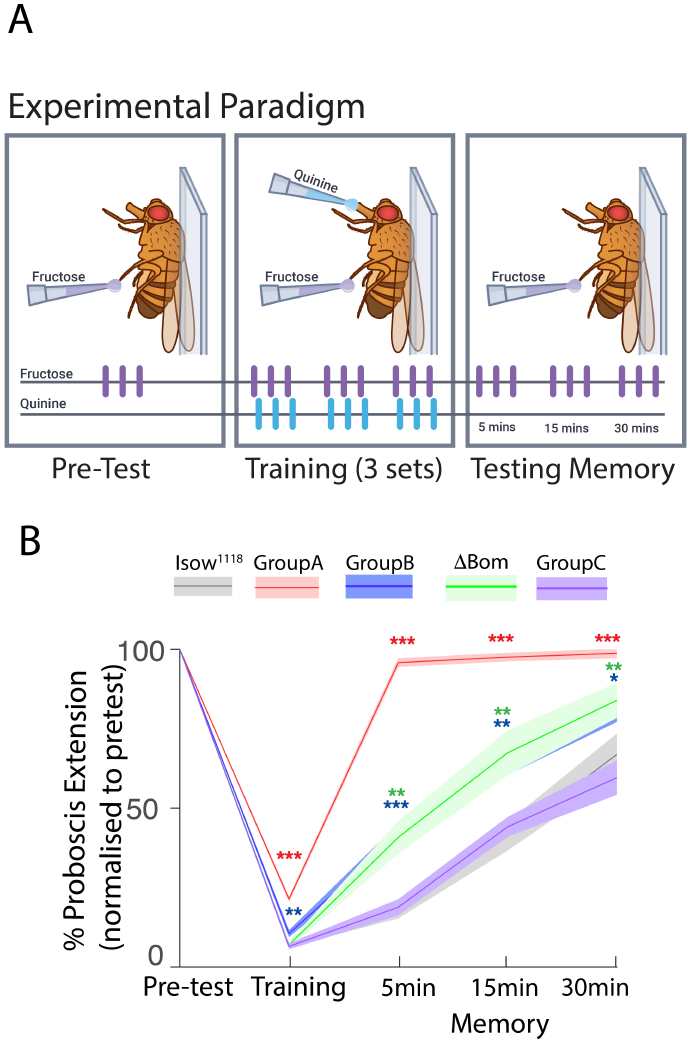
Group C mutants uniquely exhibit normal gustatory memory. (A) Schematic of the gustatory learning and memory assay. (B) Learning and memory scores (quantified as mean ± SEM proboscis extensions) of indicated genotypes are plotted. Modified Bonferroni correction following One-way ANOVA for genotype (n=11-12 flies/ genotype). * p<0.05, ** p <0.01, *** p <0.0001. ΔBom = Bomanins.

AMP mutants exhibited varying degrees of learning and memory impairments, in roughly inverse proportion to the extent of sleep loss (Fig 7B). Group A mutants showed the greatest impairments in learning and memory, while exhibiting only modest changes in sleep. Group B and Bomanin mutants which showed moderate sleep deficits also displayed moderate impairments in learning and memory. Group C mutant flies, which showed the highest sleep loss, uniquely retained normal learning and memory (Fig 7B). Importantly, none of the AMP mutants were impaired in sensory responses to fructose, sucrose, or quinine (S10 Fig).

The above data suggest the Group C mutants can carry out (some) sleep functions despite not sleeping much. Sleep loss is also known to shorten lifespan [63-66]. Accordingly, we evaluated the lifespan of AMP mutants and *iso w*^1118^ control flies, reasoning that the shortened sleep could reflect changes in lifespan. We analyzed the data using Kaplan– Meier survival analysis (S11A – S11H Fig) and mean survival comparison (S11I – S11P Fig). Group A mutants, especially males (S11A and S11E, and S11I Fig), and Group C mutants, especially females (S11D and S11H, and S11P Fig), exhibited a significantly reduced lifespan compared to control flies. Lifespan was not affected in other AMP mutants (S11 Fig), consistent with previous results [67].

### Group C genes act in glia to regulate sleep

Of all the AMP mutants, the phenotypes of Group C mutants are particularly intriguing. Group C mutants sleep the least, yet do not display many of the consequences of extended waking. How might these phenotypes be manifest?

One possibility is that the Group C mutant phenotypes are a consequence of microbiome dysregulation in these mutants. To test this hypothesis, we generated axenic Group C and *iso w*^1118^ flies and assessed their baseline sleep and sleep homeostasis. The elimination of gut microbiota had no significant effect on sleep time, sleep latency, or sleep homeostasis, in either *iso w*^1118^ controls or Group C mutant flies (S12 Fig). These results suggest that the sleep perturbations observed in Group C mutants are regulated independently of gut microbiota.

Where might the Group C genes be acting then to regulate sleep? As noted above, Group C is a double mutant in *Mtk* and *Drs*. A previous study found that overexpression of Mtk in glia increased sleep [27]. We hypothesized that *Mtk* and *Drs* function might be required in glia for sleep and sleep homeostasis. To test this hypothesis, we knocked down Mtk and Drs in glial cells using the pan-glial *repo GAL4* driver. The resulting *repo GAL4* > *UAS Drs RNAi*; *UAS Mtk RNAi, UAS Dcr2* flies recapitulated the Group C mutant sleep phenotypes, including reduced total, daytime, and nighttime sleep (S13A – S13C Fig), increased sleep latency (S13D Fig), and decreased nighttime bout length (S13F Fig) without altering waking activity (S13G Fig). Further, these flies also exhibited an exaggerated sleep rebound, like the Group C mutants (S13H and S13I Fig). These results indicate that *Mtk* and/ or *Drs* are required in glial cells to regulate sleep and sleep homeostasis.

Components of innate immunity including the Toll receptor and the NFKB transcription factors *Relish* and *Dif*, that regulate AMP expression, are known to function in the fat body and in neurons to regulate sleep [25, 26, 68].

To investigate whether Mtk and Drs are additionally required in the fat body or neurons to regulate sleep, we knocked down Mtk and Drs expression in these tissues with RNAi. We used three drivers for the fat body – *Ppl GAL4, 0*.*68 Lsp2 GAL4* and *3*.*1 Lsp2 GAL4*. Knockdown of Drs and Mtk expression in the fat body did not change sleep parameters (S14 Fig). We also used three drivers for neurons – *nsyb GAL4* (pan neuronal), *R58H05 GAL4* (R5 neurons of the ellipsoid body), and *dIlp2 GAL4* (Pars Intercerebralis). No changes in sleep time or sleep consolidation were seen upon expressing *UAS Drs RNAi; UAS Mtk RNAi, UAS Dcr2* with these neuronal drivers (S15 Fig). *R58H05-GAL4 > UAS Drs RNAi; UAS Mtk RNAi, UAS Dcr2* flies exhibited a significantly reduced latency (S15I Fig) an opposite phenotype to that observed in the Group C mutants and with RNAi knockdown in glia. The reason for this anomalous result is at present unclear. In summary, glia seems to be the major site of action of Mtk and Drs in the regulation of sleep.

### Enhancing sleep improves learning and memory in Bomanin mutants

AMP mutants exhibit sleep defects (Fig 1) and, Group C apart, show defects in learning and memory (Fig 7). Enhancing sleep has been shown to improve cognition in plasticity-impaired flies including classic mutants such as the *rutabaga* adenyl cyclase, aged flies and Alzheimer’s Disease models [58, 69-72]. We wondered whether enhancing sleep of AMP mutants could improve learning and memory. We chose to first evaluate Bomanin mutants as Bomanin mutants display moderate sleep deficits (Fig 1) and moderate memory deficits (Fig 7) and thus are a great candidate to potentially benefit from sleep enhancement. Sleep of Bomanin mutants was enhanced using two different approaches – pharmacology and behavioral medication.

To pharmacologically enhance sleep, we fed Bomanin mutants the GABA A agonist, Gaboxadol [69]. This robustly enhanced sleep (Fig 8A and S16A – S16E Fig) and improved gustatory learning and memory (Fig 8B). As a behavioral modification to enhance sleep, we used a sleep opportunity restriction paradigm [73]. In this paradigm developed by Belfer and colleagues, the sleep opportunity of flies is restricted by restricting the dark period. We switched flies from a 12:12 light: dark schedule to a 14:10 schedule, thus restricting the dark period. This intervention enhanced the sleep of Bomanin mutants, particularly at night (Fig 8C and S16H – S16L Fig), and improved memory (Fig 8D). Importantly, neither gaboxadol nor sleep opportunity restriction altered sensory responses to sugars (S16F and S16G, S16M and S16N Fig) or quinine (S10D Fig).

**Figure 8.**
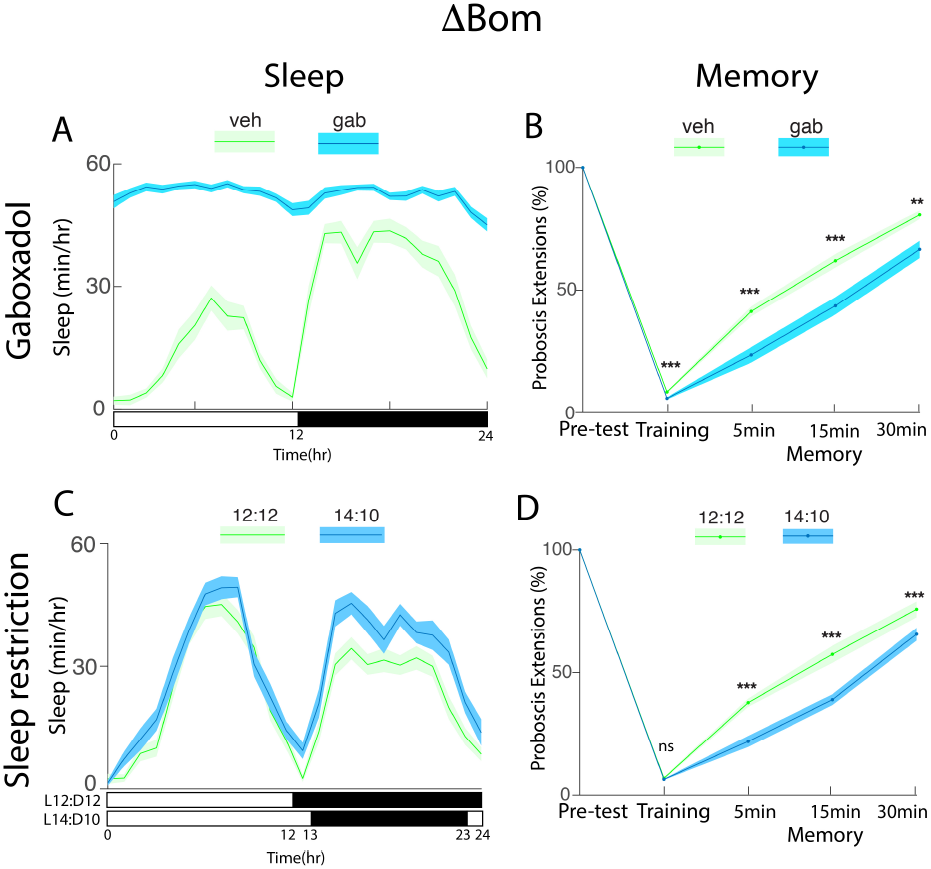
Enhancing sleep of Bomanin mutants improved gustatory learning and memory. (A) Sleep (mean ± SEM sleep) per hour for gaboxadol (gab) and vehicle (veh) fed flies as indicated. (C) Sleep (mean ± SEM sleep) per hour for flies reared in normal light-dark period (L12-D12) and prolonged light & restricted dark period (L14-D10) as indicated. The bar indicates light (white) and dark (black) period. (B & D) Learning and memory scores (quantified as mean ± SEM proboscis extensions) of indicated treatments are plotted for Bomanin mutants. (n=12 flies/genotype). ** p <0.01, *** p <0.0001. ΔBom = Bomanins.

## Discussion

Sleep and the immune response are linked. Molecules such as cytokines, and NFΚB transcription factors can, separate from their functions in host defence, regulate sleep[3, 6, 26, 74]. To better understand the role of immune molecules in the regulation of sleep, here we systematically evaluated loss of function mutants in different AMPs, effectors of the fly innate immune response. We find that sleep regulation by AMPs is common. Different classes of AMPs specifically mediated context dependent changes in sleep. Strikingly, the Group C mutants that slept the least uniquely retained (some) sleep functions. These Group C genes (Mtk & Drs) act in glia to regulate sleep and memory. AMPs thus play important roles in the modulation of sleep drive and execution of sleep functions. These results suggest a deep connection between molecular mechanisms regulating immunity and neural functions and are consistent with theories of co-evolution of immune and neural systems.

### Loss of function of AMPs affects sleep and sleep homeostasis

AMP mutants sleep less (Fig 1). Low sleep was characterised by increased sleep latency, impaired sleep consolidation, increased pWake and associated with altered arousal thresholds. Overexpression of *nemuri* and *Mtk* increased sleep [24, 27]. Based on these results, AMPs were proposed to function as somnogens. Our data extend these results with a loss of function approach to show that many classes of AMPs are required for daily sleep. Our data are also consistent with the low sleep seen in mutants in the NFΚB transcription factors *Dif* and *Relish* [25, 26].

Expression of many AMPs, including *Mtk* (one of the Group C mutants), was shown to be elevated following sleep deprivation[27]. We next evaluated sleep homeostasis, reasoning that low sleep might result from an inability to generate needed sleep. Sleep rebound was unaltered in most AMP mutants; Group C mutants uniquely exhibited an increased rebound. A close examination of rebound dynamics, however, revealed an interesting pattern. Group B and Group C mutants were impaired in the extent of sleep rebound after a 12h recovery period, but caught up, and in the case of Group C, exceeded the wild-type rebound after 48h of recovery. These results reinforce the value of evaluating rebound over a 48-hour period. Group C mutants also recovered a substantial fraction of lost sleep at night, in contrast to wild-type flies, where the clock is known to gate the period of recovery such that sleep is recovered during the day. Our data suggest that the sleep homeostat is perturbed in interesting ways in Group C mutants and form the basis of more mechanistic studies to follow.

The two process model of sleep regulation posits that sleep is regulated by circadian and homeostatic processes. Accordingly, we examined circadian behaviour in AMP mutants. Group C mutants showed a mild shortening of the circadian period, as well as lowered morning and evening anticipation. A microarray analysis of clock regulated genes identified many AMPs including the Group C genes *mtk* and *drs*, and the Group B genes *Attacin A (Att A), Attacin B (Att B)* and *drosocin (dro*) [75]. Further, an analysis of the cycling transcriptome in astrocytes showed that expression of one of the Bomanin genes, BomBc2 cycles in astrocytes [76]. Our data are consistent with these expression profiling experiments and show that AMPs are functionally required for aspects of circadian behaviour.

### Specific AMPs mediate context dependent changes in sleep

AMP mutants affect baseline sleep. Sleep is also however known to be plastic and modifiable by environmental changes and ecological niches. To determine if, in addition to regulating baseline sleep, AMPs also mediate sleep plastic changes in sleep, we first evaluated socialization induced changes in sleep. Flies housed in groups sleep more following the period of socialization than their isolated siblings[41]. Group B and Bomanin mutants were impaired in socialization induced sleep. Of note, expression of a number of AMPs including the Group B genes *AttB* and *dro* were shown to be modulated by social experience [77]. Group B and Bomanin mutants were also impaired in sleep induced by gentle rocking (Fig 4B). Little is hitherto known about the molecular mechanisms that mediate rocking induced sleep. Disrupting flight of flies is known to induce long lasting changes in sleep [50]. We find that flight-disruption induced sleep is unaltered in AMP mutants. These results are consistent with previous findings that flight-disruption induced sleep in flies mutant for *imd* or *relish* is unaltered [50].

The above results are examples of sleep-plastic increases in sleep. Sleep can also be reduced in different circumstances. Starvation reduces sleep in animals from flies to humans[46, 47, 53, 78]. AMP mutants exhibited sex specific changes in starvation induced sleep suppression. Group C mutant females were impaired in starvation responses, Bomanin mutant females showed an exaggerated response. The GroupC and Bomanin genes have not been implicated in starvation induced sleep suppression. A recent study showed that the Group B genes *Att B* and *Attacin C* (*Att C*) were required in a specific subset of Leukokinin expressing neurons for starvation induced sleep suppression [79]. We find that GroupB mutants show changes in starvation induced sleep suppression when day and night sleep are considered separately (S8 Fig). Different Group B mutants might interact such that individual mutants affect aspects of starvation sleep suppression but together no effect is seen on total sleep. We also examined the rebound sleep of AMP mutants following a day’s starvation. Group C mutants displayed an exaggerated rebound i.e. they recovered more sleep than controls (despite losing less sleep than controls) (S8 Fig). This is consistent with the exaggerated rebound seen in Group C mutants following overnight sleep deprivation. GroupA mutants also displayed an exaggerated rebound following starvation while being unaffected in response to mechanical sleep deprivation. These results reveal a context dependent role for AMPs in sleep regulation. Together, these data suggest that while AMP regulation of sleep might be common, different AMPs regulate particular aspects of context dependent changes in sleep.

### Group C mutants retain some sleep functions

Our data are thorough characterisation of the role of AMPs in sleep regulation. We next evaluated sleep functions. The Synaptic Homeostasis Hypothesis (SHY) is an influential theory of sleep function that posits that waking experience upscales synapses which are pared back during sleep to facilitate encoding of new information the next day [54]. Consistent with the predictions of SHY, extended waking induced by sleep deprivation or in various sleep short sleep mutants resulted in elevated levels of the presynaptic protein Bruchpilot (BRP) [55, 56, 80, 81]. Upon examining AMP mutants, we found that the brains of Group C mutants uniquely displayed lowered synapse abundance as evidenced by lowered BRP levels. This result is surprising on multiple levels. Group C mutants sleep the least, hence, as per SHY, one would expect elevated BRP levels. Further, elevated BRP levels following extended waking is thought to drive sleep need. Yet, Group C mutants display lower BRP levels, and an enhanced sleep rebound. A deeper mechanistic understanding of these phenotypes awaits a thorough biochemical characterization of AMP function. In this regard, it is worth noting that the immediate early gene Homer is thought to mediate sleep dependent synaptic scaling [82], Homer mutants in flies also displayed altered sleep homeostasis [83]. Potential interactions between Homer and the Group C peptides Mtk and Drs are certainly worth exploring. Interestingly, in mammals, the Major Histocompatibility Complex 1 and complement proteins have been shown to be involved in synaptic pruning [84, 85].

From worms to humans, sleep and sleep loss are known to be important for memory consolidation [86]. Accordingly, we evaluated learning and memory in AMP mutants using a well established gustatory learning and memory paradigm [62]. Learning and memory in AMP mutants was in roughly inverse proportion to the extent of sleep loss (Fig 7). Group C mutants uniquely retained normal learning and memory despite sleeping the least. Taken together with the previous result these data suggest that Group C mutants are able to carry out (some) sleep functions despite not sleeping very much. Group B mutants displayed learning and memory impairments in our assay. A recent study from Barajas-Ezpeleta and colleagues evaluated long term memory following knockdown of different AMPs in neurons and in the fat body [87]. Bomanins and Mtk were not evaluated in this study, which identified the Group B gene *DptB* as being required for long term but not short term memory. These results can be reconciled with our data by considering a few possibilities. First, one of Att A, Att B, Att C, Dpt A or Dro could be uniquely required for short-term but not long-term memory. Second, Att A / Att B / Att C/ Dpt A or Dro could act in glia to modulate memory (short and long-term). Third, mutants in Group B genes could in combination impair short term memory. Evaluating single mutants in different Group B genes would help distinguish between these possibilities. Interestingly, a short peptide *Induced by Immunity (IBIN)* was shown to be required for the ability of flies to modulate their mating behavior is response to parasitoid wasps, a form of non associative learning [88]. Expression of IBIN was shown to be upregulated in response to bacterial infection in immune tissues, it could also be activated by either the Toll or *imd* pathway, indicating that it is likely an immune peptide [89].

Our finding that Group C mutants can retain some sleep functions despite their low sleep are exciting. While a vast body of literature supports the critical role for sleep in cognition and optimal functioning, a growing body of work suggests that animals can maintain functioning without much sleep in response to environmental challenges or ecological niches. A study of polygynous pectoral sandpipers found that they maintain high behavioural performance over several weeks despite greatly reduced sleep during this period [90]. Interestingly, in this study, the males that slept the least appear to sire the most offspring [90]. Similarly, Rattenborg and colleagues found that white-crowned sparrows during the migratory season maintained a high level of cognitive performance on a repeated-acquisition task despite sleeping abut 2/3rds of the duration in the non migratory season[91]. Several strains of blind Mexican cavefish *Astyanax mexicanus* sleep very little [78]. Further, dolphins and killer-whales mothers and calves were shown to suppress sleep for weeks following birth as they migrate from one site to another[92]. A study that examined elephant sleep in natural conditions, found that elephant sleep little in the wild, and could on occasion go days without sleep [93]. Our data with the Group C mutants add to this body of work, and hint at potential mechanisms that might enable animals to maintain functioning during low sleep.

### Sites and mechanisms of action of AMPs

Where might these genes be acting to influence sleep and memory? To answer this question, we turned to RNAi mediated knockdown. We focussed on the Group C genes as we observed the strongest effects with Group C mutants, and it is tractable to RNAi based knockdown. The NFΚB *Relish* regulates sleep from the fat body [25]. A different NFΚB, *Dif* regulates sleep in neurons including the Pars Intercerebralis [26]. Knockdown of Drs and Mtk using drivers that express in neurons, or the fat body did not change sleep (S14 Fig). In contrast, knockdown of Drs and Mtk in glia resulted in low sleep, increased latency, and an exaggerated sleep rebound. The sleep phenotypes of Mtk & Drs loss of function thus map to glia. Glial expression of AMPs might be common. BomBc2, one of the genes deleted in the Bom^Δ55c^ allele we used, was shown to be expressed in astrocytes and regulate sleep [76]. BaraA and Bara C, two Baramicin peptides are also expressed in glia [23, 94]. Indeed, BaraA’s functions in immunity also mapped to glia – BaraA secretion from glia was shown to neutralize fungal toxins[95]. Knockdown of the NFΚB transcription factors Dif and Relish in glia did not change sleep [26]. Mtk and Drs expression in glia thus maybe regulated by *dorsal* or by FOXO transcription factors [96]. Interestingly, in addition to glia, at least one of the Group C genes - Mtk, is expressed in and functions in neurons. Knockdown of Mtk in neurons was neuroprotective in an amyotrophic lateral sclerosis model [97].

How might these genes be acting to regulate nervous system functions? AMPs are classically thought to act as membrane disruptors, binding to negatively charged lipids by virtue of their positive charge [98]. However, it is increasingly clear that AMPs can act on intracellular targets as well. In a heterologous system, Mtk bound to and inhibited succinate coenzyme Q reductase on mitochondria [99]. Drs was shown to modulate the action of a toxin on a voltage gated sodium channel [100]. Dro, a proline rich AMP, was shown to bind to the ribosome and inhibit translation [101]. While these studies have not addressed potential mechanisms by which AMPs might serve non immune functions, the diversity of targets throw up many possibilities. In *C. elegans*, the AMP NLP29 was shown to bind to a GPCR NPR12 and regulate dendrite degeneration and sleep [102, 103]. While AMPs were first identified for their antimicrobial properties, the extent to which they function as antimicrobials vs (neuro) peptides remains unresolved and is an area of active investigation.

## Conclusion

These data are a thorough, systematic characterization of the roles of AMPs in sleep regulation and function. We have taken a comparative approach that allows us to discover general themes of AMP action in sleep, as well as specific roles for particular classes of AMPS. That many classes of AMPs regulate sleep suggest that neural and immune systems may have co-evolved, with immune genes being recruited to serve neural functions. Here, as a first pass, we examined AMP mutants in different groups – Group A (mutant for 3 cecropins + Defensin), Group B (mutant for 4 Attacins, 2 Diptericins, and Drosocin), Group C (mutant for Mtk and Drs), and Bomanins (a deletion in 10 Bomanin genes). Going forward, examining single mutants in different AMP genes might yield a more nuanced understanding of the role of specific AMPs. The Group C mutants are particularly intriguing as they appear to be able to carry out (some) sleep functions sans much sleep. We expect that future investigations into the specific roles of Mtk & Drs will yield important broadly applicable insights into mechanisms of sleep regulation and function.

## Materials and Methods

### Fly strains and handling

Flies were reared on Bloomington recipe standard cornmeal agar food and maintained at 25°C with approximately 50-55% relative humidity and a 12-hour light:12-hour dark schedule. Antimicrobial peptide (AMP) mutant strains-Group-A, Group-B, Group-C, ΔBomanin, and *iso w*^1118^ controls were a kind gift from Bruno Lemaitre (EPFL, Switzerland).

The following lines were obtained from the Bloomington Drosophila Stock Center: *Da GAL4*; *Da GAL4* (RRID:BDSC_95282), *Ppl GAL4* (RRID:BDSC_58768), 0.68*Lsp2 Gal4* (RRID:BDSC_6357), 3.1*Lsp2 GAL4* (RRID:BDSC_98128), *nSyb GAL4* (RRID:BDSC_51941), *dIlp2 GAL4* (RRID:BDSC_37516), *R58H05 GAL4* (RRID:BDSC_39198), *repo GAL4* / Tm3Sb (RRID:BDSC_7415), *UAS Mtk RNAi*/ TM3Sb (RRID: BDSC_28546), *UAS Drs RNAi* (RRID:BDSC_63631), *UAS Dcr2* (RRID:BDSC_24651).

### Sleep Behaviour

Sleep was measured using Drosophila Activity Monitors (DAM2, Trikinetics Inc., Waltham, MA, USA). Newly eclosed flies were individually housed in 65 mm glass tubes containing standard cornmeal agar food at one end and the other end stoppered with foam. The glass tubes were placed into the activity monitors, and their locomotor activity was continuously monitored. The locomotor activity was binned in 1-minute intervals, and sleep was defined as the period of inactivity lasting five minutes or longer. Sleep duration was computed using custom macro-enabled Excel scripts [65]. Baseline sleep was evaluated in 4 to 6-day-old flies maintained in a 12-hour light and 12-hour dark schedule.

### Arousal Threshold

Arousal Thresholds were measured using the *Drosophila* Arousal Tracking (DART) software [34] in a custom-built apparatus. Non-mated female flies (5-6 days old) were individually placed in 75 mm long glass tubes. 16 flies per group were loaded in tubes placed in a 3D-printed platform. Three vibration motors (Precision Microdrives) were glued under each platform. Flies were illuminated with infra-red light (825nm, LED Lights World). Fly activity was video monitored continuously at 5 frames per second for 2 days using a USB webcam (Model no. c920, Logitech, Lausanne, Switzerland) with the infrared filter removed. The DART software interface was used to deliver increasing voltage pulses to the motors via analog output channels of a USB data acquisition device (Model no. 1280 LS, Measurement Computing, Newbury, Berkshire, UK).

### Sleep Homeostasis

Sleep homeostasis was evaluated using established protocols [36]. Virgin female flies (4-5 days old) were placed in DAM tubes, and their baseline sleep was recorded over 2 days. Flies were then deprived of sleep for 12 hours during the dark phase (ZT12-ZT0) by placing the DAM monitors on the Sleep Nullifying Apparatus (SNAP). After deprivation, the flies were left undisturbed for two days to monitor their sleep rebound. For each fly, the difference in sleep duration on recovery days 1 and 2 and the baseline day was calculated as the sleep gained or lost. Further, the percent rebound was calculated for individual flies as the percentage of the sleep gained or lost during recovery relative to the baseline day.

### Circadian Analysis

Male flies aged 4-5 days were monitored for baseline activity patterns for four days under a 12-hour light and 12-hour dark (LD) cycle. The monitors were then placed in a custom-made light-proof black box to maintain constant darkness for eight days (DD), allowing free-running activity patterns to be recorded in the absence of light cues. Circadian analysis was performed using the SleepMat software [104] on locomotor activity data collected through the Drosophila activity monitors.

### Social Enrichment

Socialization induced sleep was evaluated following established protocols [41]. Newly eclosed flies were collected and housed in groups of 40-42 flies per vial for five days to allow social enrichment. In parallel, a control group of isolated flies was individually collected and transferred to 65mm DAM tubes and kept for five days. After five days of social enrichment in vials, the enriched flies were loaded into individual 65mm DAM tubes. The socialization-induced change in daytime sleep was calculated as ΔSleep [enriched-isolated] for each genotype. The change in sleep of AMP mutants was then expressed as a percent of the change in sleep of *iso w*^1118^ controls.

### Rocking-Induced Sleep

The orbital rocking protocol was adapted from Lone et al [45]. Virgin female flies (5-7 days old) were loaded in *Drosophila* activity monitors (Trikinetics Inc., Waltham, MA, USA). These monitors were securely attached with masking tape to the surface of a laboratory orbital rocker (Catalog #88882008, ThermoFisher Scientific, Waltham, MA, USA) operating at 90 rpm during the light phase (ZT1-ZT12). Baseline sleep was monitored for 3 days post-eclosion, and then the flies were placed on the rocker for 3 days to undergo daytime rocking. The average change in sleep over 3 days of rocking was compared to the baseline sleep to evaluate the change in sleep during the rocking phase.

### Starvation-Recovery Assay

Sleep of 4 to 5-day-old flies was monitored for two days on regular food using the DAM system. This served to establish a baseline for sleep. Once sleep was determined to be stable, flies were transferred to tubes containing 1% agarose for 24 hours of starvation during which sleep was monitored. Following 24h of starvation, flies were returned to standard food for 24 hours, during which sleep was monitored as recovery sleep. The difference in sleep between starvation day and baseline, and recovery day and baseline, was then computed for each genotype. This change in sleep was expressed as the percent sleep lost during starvation and regained during recovery.

### Wing cut

Newly eclosed female flies were collected, and both wings were cut following established procedures using micro scissors under CO2 anesthesia [50]. Flies with cut wings and intact wing controls that had been subjected to the same CO_2_ anesthesia were then transferred to 65-mm glass tubes and loaded onto DAM monitors to record sleep. Sleep data was recorded for three days after the wing cut. The change in total sleep time cut-intact was computed for each genotype tested. The change in sleep of AMP mutants was then expressed as a percent of the change in sleep of *iso w*^1118^ controls.

### Immunohistochemistry

Adult male flies aged 7-9 days were anesthetized on ice and fixed in 4% paraformaldehyde (diluted from 20% stock, Catalog #15713, Electron Microscopy Sciences, Hatfield, PA, USA) in 1X Phosphate Buffered Saline (Catalog # P3813, Merck, St. Louis, MO, USA) with 0.4% Triton (Catalog # X-100, Merck, St. Louis, MO, USA) (PBST) for 2 hours. The animals’ brains were then dissected in PBS and collected in PBST on ice. Brain tissues were washed 3 times (15-20 min each) with cold PBST and blocked using 5% goat serum (Catalog # RM10701, HiMedia, Mumbai, IN) in PBST for 3 hours at room temperature (25°C) and then incubated with primary antibodies diluted in 5% goat serum in PBST for 48 hours at 4°C with gentle rocking. The samples were then washed 3 times with PBST and incubated with secondary antibodies in 5% goat serum/PBST at 4°C for 24 hours with gentle rocking in the dark. After the secondary antibody staining, the brain samples were washed 3 times with ice-cold PBST and then transferred into PBS. The whole brain samples were mounted in VectaShield reagent (Catalog #H-1000, Vector Labs, Newark, CA, USA) on poly-L-lysine-coated glass slides (Catalog #, Electron Microscopy Sciences, Hatfield, PA, USA) and imaged using a Zeiss LSM900 confocal microscope. Primary antibody used: mouse anti-BRP (mAb-nc82, Developmental Studies Hybridoma Bank, Iowa City, IA, USA) at 1:400. Secondary antibody: Alexa Fluor 568 goat anti-mouse IgG (RRID:AB_2534072, ThermoFisher Scientific, Waltham, MA, USA) at 1:200.

### Gustatory Learning Assay

Mated female flies (7 to 9 days old) were used to assess gustatory learning and memory using a gustatory associative conditioning assay following established procedures [62]. Flies were wet-starved in a vial with water-soaked Kimwipe overnight before the experiment. The following morning, they were thorax-tethered to glass slides using UV-activated glue (Laser Bonding Tech-BondicModel: SK8024) and allowed to acclimate for 45 minutes. Baseline proboscis extension response to fructose solution (100 mM, Catalog #F0127, Merck, St. Louis, MO, USA) applied on their tarsi was counted for three trials of 10 seconds each. The training consisted of pairing fructose presentation to the tarsi with quinine (10 mM, Catalog #Q1125, Merck, St. Louis, MO, USA) to the proboscis upon extension, repeated over three sets of three trials each. The inter-trial interval was 10 seconds, and the inter-set interval was 3 minutes. Memory was assessed after 5-, 15-, and 30-minute post-training by measuring the proboscis extension response to fructose presentation on the tarsi alone. All gustatory learning assays were conducted between ZT1 and ZT5. Data were normalized to baseline response and analyzed using appropriate statistical tests as indicated.

### Quinine Sensitivity Assay

Quinine taste sensitivity was assessed using a choice assay between a 50 mM fructose solution and a mixture of 50 mM fructose + 5 mM quinine. Mated female flies aged 7-9 days, reared on standard medium, were wet-starved for 24 hours in a vial with water-soaked Kimwipe before the assay. Groups of 5-6 flies were aspirated into a 35 mm petri dish with a lid. In the Petri dish, flies were presented with a choice between a drop of quinine + fructose and fructose-only solutions. The solutions were colored with neutral food coloring dyes (blue or red) to visually track consumption. Dye colors were swapped between trials such that each genotype was tested with both combinations. Flies were given 30 minutes to perform the choice assay in a group of 6-8 flies per trial. Following the assay, flies’ abdomens and proboscises were inspected under a microscope to identify which solution had been consumed or if flies had consumed both solutions. The number of flies choosing each option was counted to evaluate their ability to detect and avoid quinine.

### Gaboxadol feeding

Gaboxadol (Catalog #T101, Merck, St. Louis, MO, USA) was fed to flies mixed in the fly food at a concentration of 0.1 mg/mL following a previously described protocol [69]. Female flies were fed Gaboxadol mixed food for two days before gustatory learning and memory testing.

### Sleep Opportunity Restriction

The sleep opportunity restriction protocol was adapted from Belfer et al. [73]. Newly eclosed female flies were transferred into 65mm glass tubes and their baseline sleep was monitored inside a 25°C incubator with a 12-hour light and 12-hour dark (LD) cycle for 3-4 days. Following this, the flies were shifted to a restricted dark period of 10 hours using a controlled 14-hour light and 10-hour dark cycle in the incubator for 4-6 days.

### Life Span Assay

Newly eclosed flies were collected and transferred into vials housed in groups of 8 per vial with standard fly food. 8–9 such vials of males and females were set up for each genotype. The flies were transferred to fresh food vials every three days to ensure fresh food and optimal conditions for the flies. The lifespan duration of each fly was manually observed and recorded in days. The lifespan data were analyzed for AMP mutants and the control-*iso w*^1118^ flies using the log-rank test and comparing the mean survival of flies in each vial for statistical significance. Kaplan-Meier survival curves were generated using custom Python scripts.

### Axenic flies

Axenic flies were generated following established protocols [105, 106]. Adult flies were allowed to lay eggs on fresh food plates for 2-3 hours. Embryos aged 14-16 hours were collected and dechorionated using 2.7% sodium hypochlorite solution (2-fold diluted bleach) for 3-4 minutes. The embryos were then rinsed twice with 70% ethanol, followed by 3 washes with distilled, sterile water to remove any residual bleach and ethanol. Processed embryos were transferred under sterile conditions into autoclaved axenic food vials. Handled embryos (controls) underwent the same procedure but were treated only with sterile distilled water. Axenic status was confirmed by plating homogenates from 4–5 adult flies on LB agar to check for microbial growth.

### Statistical analysis

Sleep time data are plotted as the average along with the standard error of the mean (SEM) in all sleep line plots (e.g., in Figure 1A). Most sleep characteristics are plotted as scatter plots with a line representing the mean and raw data points. Arousal threshold data were non-normally distributed and are presented as a violin plot. Graphs were plotted in MATLAB using custom scripts (Natick, MA, USA), and figures were put together in Adobe Illustrator (Adobe Inc., San Jose, CA, USA). Statistical analyses were done using GraphPad Prism software (GraphPad Software, Inc., La Jolla, CA, USA). Statistical comparisons between two groups were done with a student’s t-test or Mann-Whitney U test, where appropriate. Comparisons between more than two groups were done with a one-way ANOVA followed by a modified Bonferroni test unless otherwise noted. Kruskal-Wallis ANOVA followed by Dunn’s post-hoc test was used for all non-normally distributed data. Statistical significance was defined as p < 0.05.

## Acknowledgements

The authors gratefully acknowledge the support of the Ashoka University Microscopy Facility, where the confocal images in Fig 6 were acquired. The authors are also grateful to Prof Paul Shaw for critical comments on the manuscript and to Sumita Nanda for assistance with schematics. Research in KM’s lab is supported by Ashoka University, a Core Research Grant from the Science and Engineering Research Board, Govt of India (CRG/2022/006967), and a Ramalingaswami Re-entry Fellowship from the Dept of Biotechnology, Govt of India.

This paper was typeset with the bioRxiv word template by @Chrelli: www.github.com/chrelli/bioRxiv-word-template

## Author contributions

**Conceptualization**: Imroze Khan and Krishna Melnattur

**Data curation**: Rahul Kumar, Gokul Madhav, Krishna Melnattur

**Formal analysis**: Rahul Kumar and Krishna Melnattur

**Funding acquisition**: Imroze Khan and Krishna Melnattur

**Investigation**: Rahul Kumar, Gokul Madhav, Priyanka Balasubramanian, Rhea Lakhiani, Mugdha Joshi, Vibhu Jaggi, Anushna Pal and Krishna Melnattur

**Methodology**: Rahul Kumar, Gokul Madhav Krishna Melnattur

**Project administration**: Rahul Kumar and Krishna Melnattur

**Resources**: Krishna Melnattur

**Software**: Rahul Kumar and Krishna Melnattur

**Supervision**: Rahul Kumar and Krishna Melnattur

**Validation**: Krishna Melnattur

**Visualization**: Rahul Kumar and Krishna Melnattur

**Writing**: original draft-Rahul Kumar and Krishna Melnattur

**Writing**: review & editing-Rahul Kumar and Krishna Melnattur

