## Supplementary Figs for "Mutations in antimicrobial peptides differently affect sleep and plasticity"

#### Supplementary Figure 1

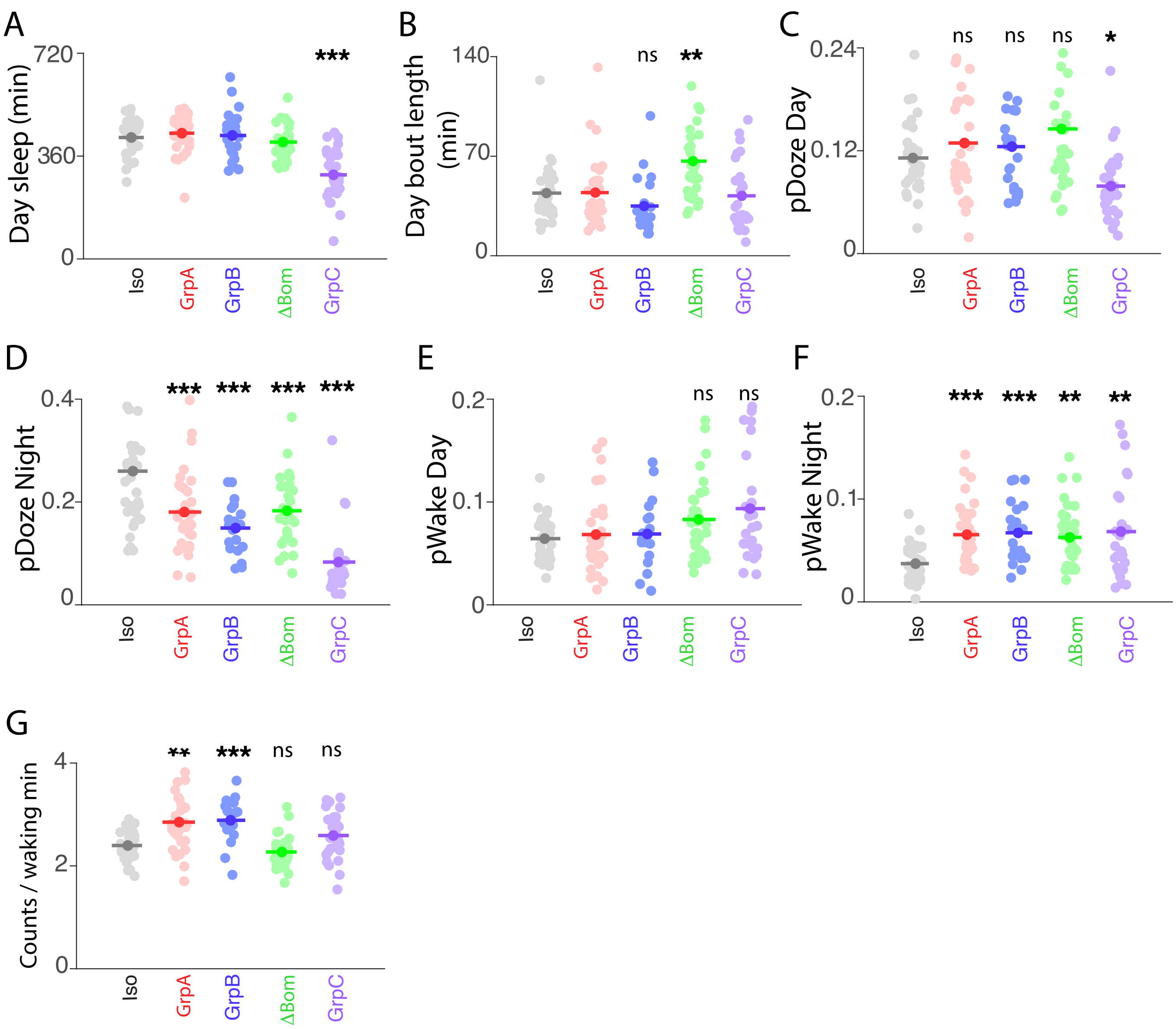


**Supplementary Fig1- Additional sleep characteristics of male AMP mutants.** Mean and raw data points are shown for (A) day sleep, (B) day bout length, (C) pDoze Day, (D) pDoze night, (E) pWake day, (F) pWake night, and (G) waking activity. Sleep characteristics of mutants were compared to *iso w*[1118] controls. (A & D) Modified Bonferroni correction following one way ANOVA for genotype. (B & C, E - G) Dunn's multiple comparisons following Kruskal-Wallis ANOVA (n=24-32 flies/genotype). * p<0.05, ** p <0.01, *** p <0.0001. Iso = *iso w*[1118], GrpA = Group A, GrpB = Group B, ΔBom = Bomanins, GrpC = Group C.

### Supplementary Figure 2

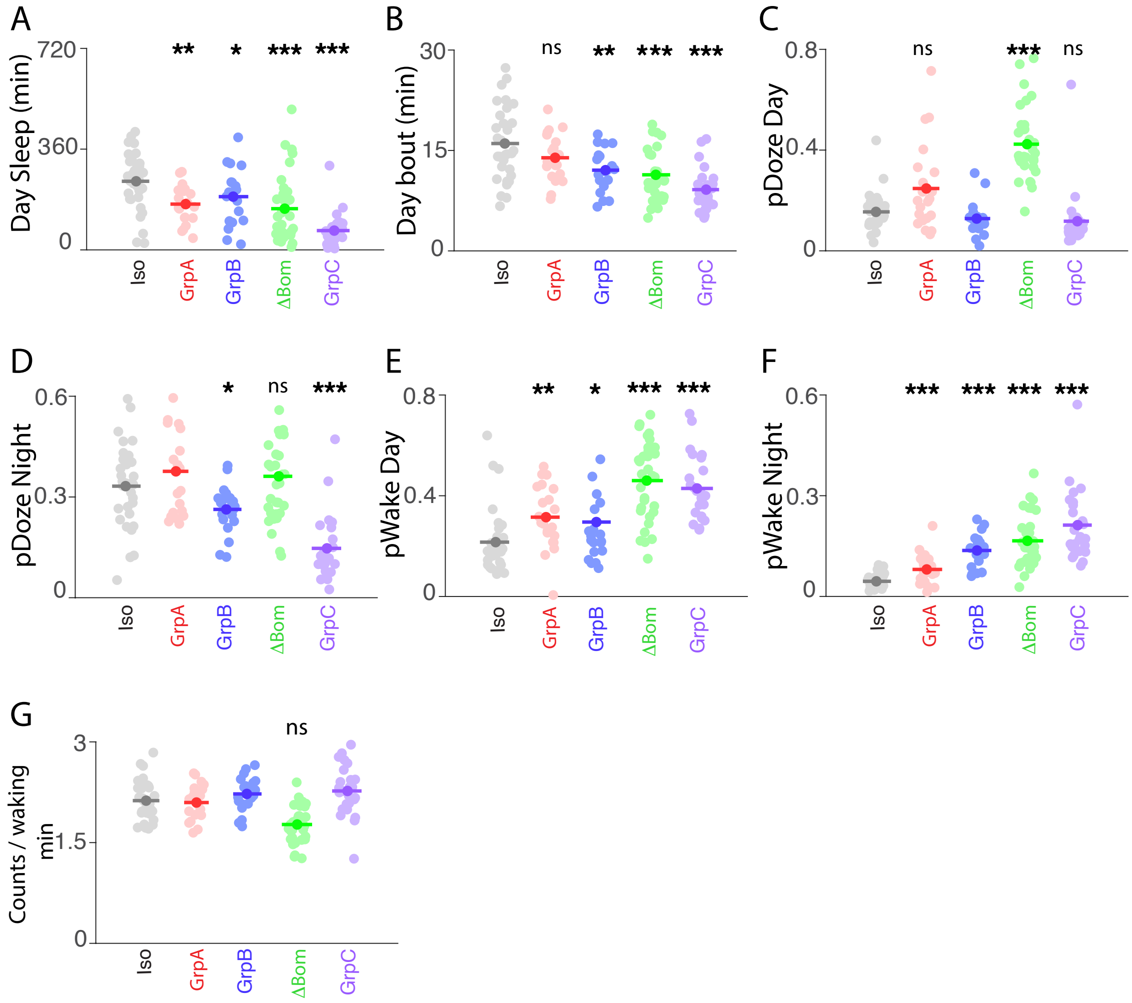


**Supplementary Fig2- Additional sleep characteristics of female AMP mutants.** Mean and raw data points are shown for (A) day sleep, (B) day bout length, (C) pDoze Day, (D) pDoze night, (E) pWake day, (F) pWake night, and (G) waking activity. Sleep characteristics of mutants were compared to *iso* $w^{1118}$ controls. (A & B, D - G) Modified Bonferroni correction following One way ANOVA for genotype. (C) Dunn's multiple comparisons following Kruskal-Wallis ANOVA (n=24-32 flies/genotype). * p<0.05, ** p <0.01, *** p <0.0001. Iso = *iso* $w^{1118}$, GrpA= Group A, GrpB = Group B, ∆Bom = Bomanins, GrpC = Group C.

### Supplementary Figure 3

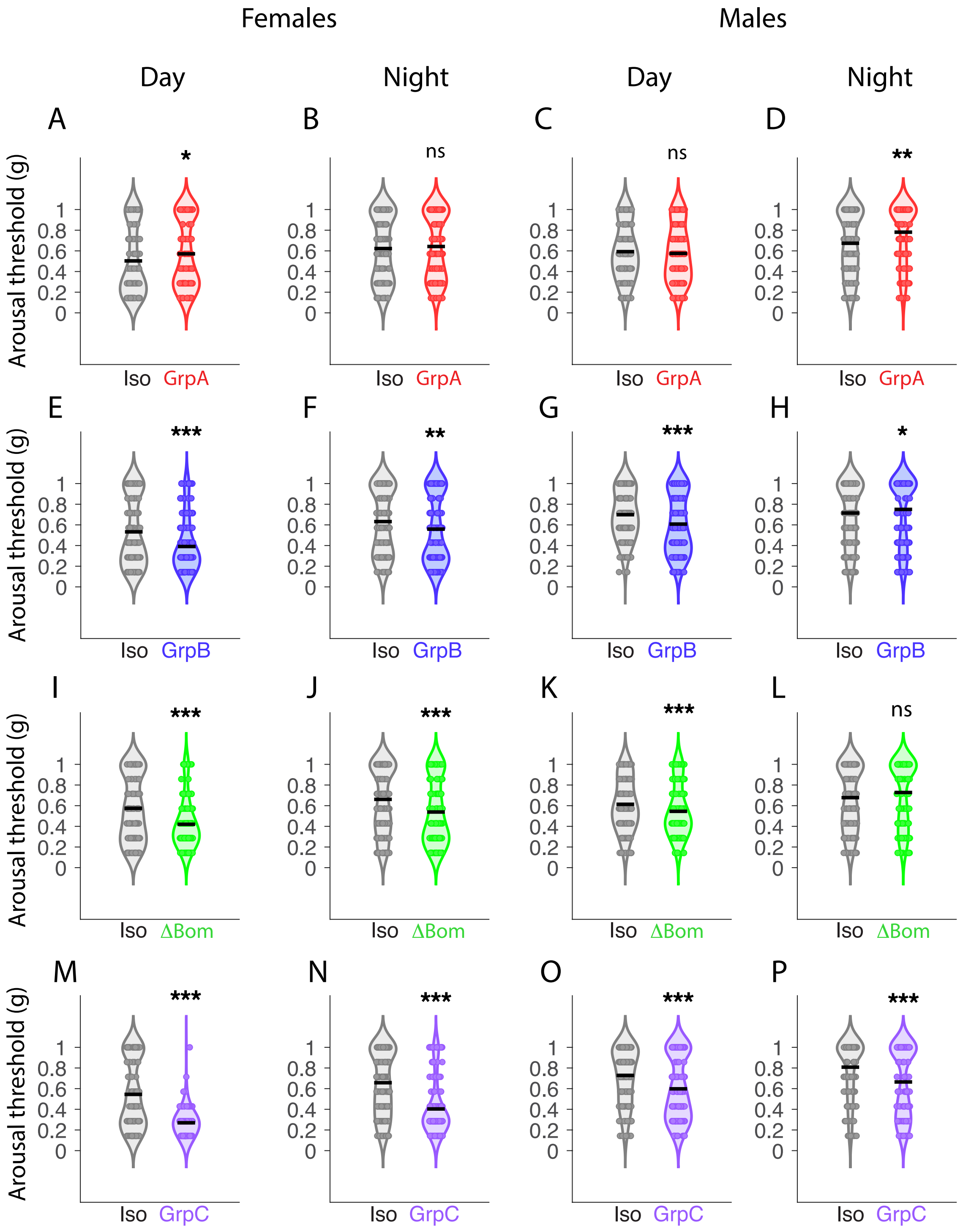


**Supplementary Fig3- Arousal thresholds of AMP mutants.** (A- P) Violin plot with mean and raw data points for arousal events at different stimulus intensities for *Iso* $w^{1118}$ controls and mutants. (A, E, I, M) females day arousal, (B, F, J, N) females night arousal, (C, G, K, O) males day arousal, (D, H, L, P) males night arousal. Mann-Whitney, one-tailed test (n=16 flies/genotype). * p<0.05, ** p <0.01, *** p <0.0001. Iso = *iso* $w^{1118}$, GrpA = Group A, GrpB = Group B, ΔBom = Bomanins, GrpC = Group C

#### Supplementary Figure 4

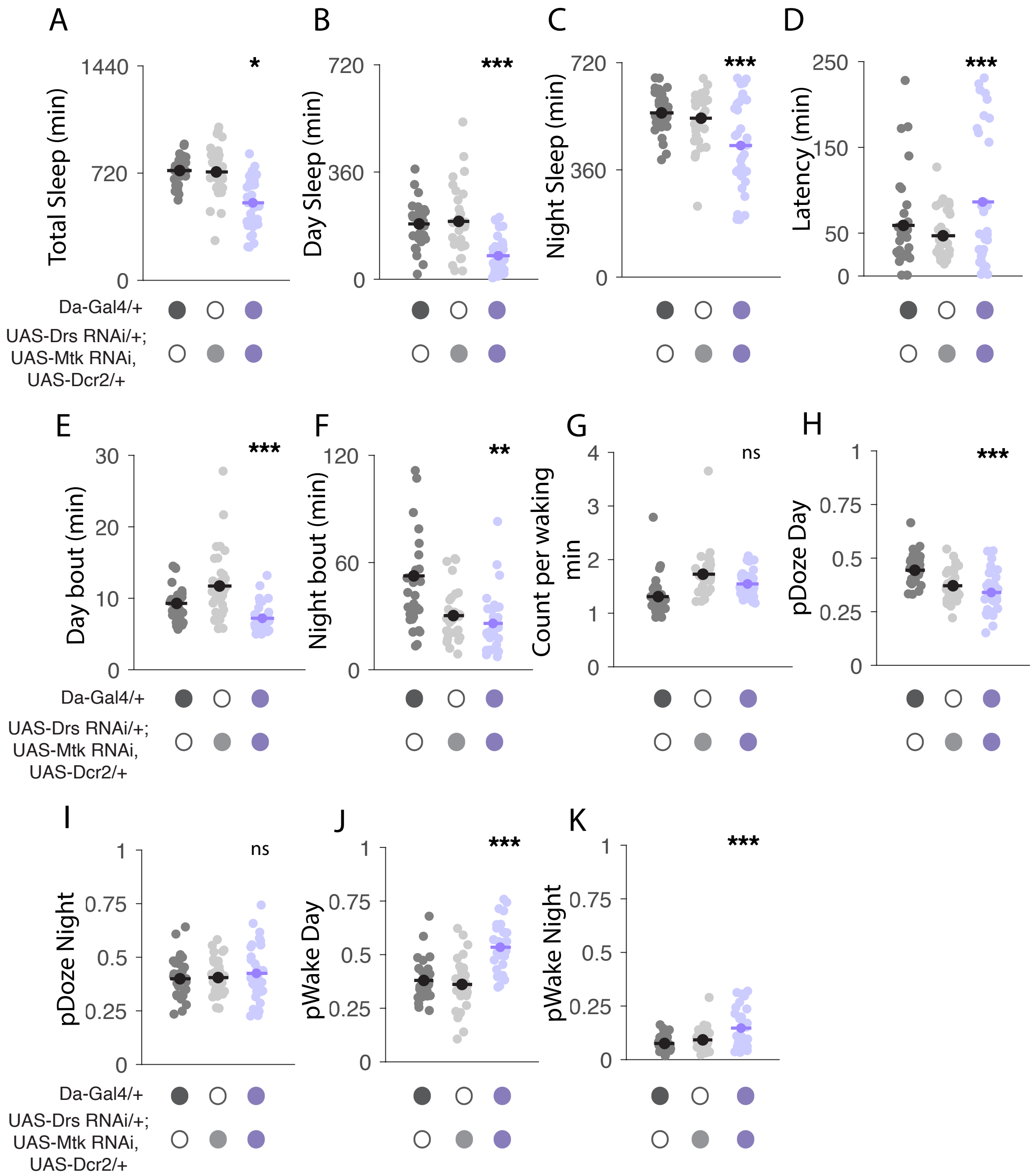


**Supplementary Fig4- Ubiquitous knockdown of Metchnikowin and Drosomycin reduced sleep.** Mean and raw data points are shown for (A) total sleep, (B) day sleep, (C) night sleep, (D) latency, (E) day bout length, (F) night bout length, (G) waking activity, (H) pDoze day, (I) pDoze night, (J) pWake day, (K) pWake night. Sleep characteristics of *Da GAL4> UAS-Drs RNAi, UAS Mtk RNAi, UAS Dcr2* were compared to their genetic controls- *Da GAL4*/+ and *UAS Mtk RNAi, UAS Dcr 2* / +, *UAS-Drs RNAi* /+. (A-K) Modified Bonferroni correction following one way ANOVA for genotype (n=31-32 flies/genotype). *p<0.05, **p <0.01, ***p <0.0001.

#### Supplementary Figure 5

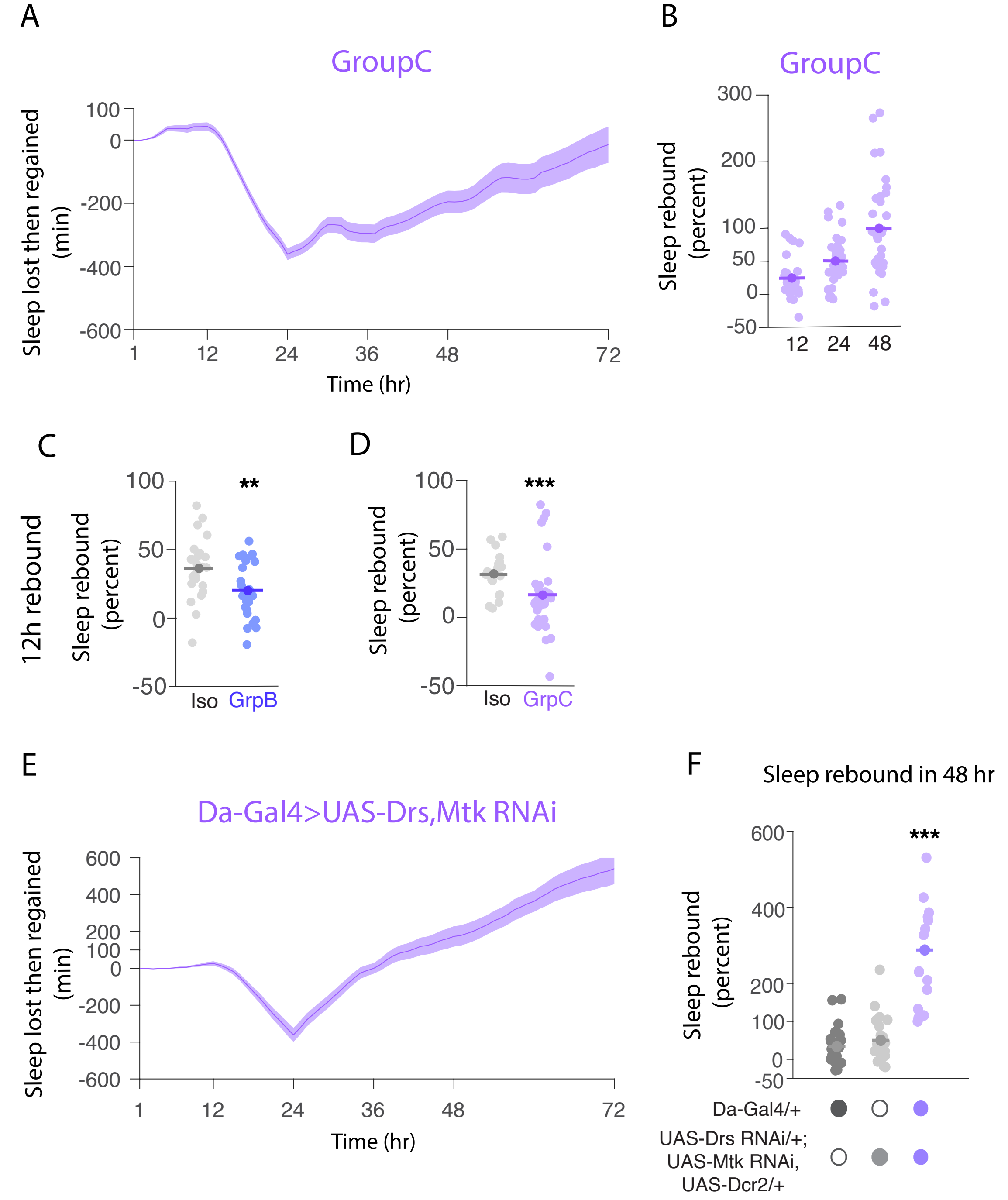


**Supplementary Fig5- Rebound sleep characteristics of Group C null mutant and knockdown animals.** (A & E) Plot of mean ± SEM sleep lost during overnight sleep deprivation and gained during 48h of recovery for Group C (A) and *Da GAL4> UAS-Drs RNAi*, *UAS-Mtk RNAi*, *UAS Dcr2* (E) is shown. (B) Percent sleep rebound for Group C mutants over 12h, 24h, and 48h post sleep deprivation. (C & D) Percent sleep rebound over 12h for *iso* $w^{1118}$ controls compared to Group B (C) and Group C (D), respectively. (F) Percent sleep rebound over 48h for *Da GAL4> UAS-Drs RNAi*, *UAS-Mtk RNAi*, *UAS Dcr2* and its genetic controls – *Da GAL4*/+ and *UAS Mtk RNA*i, *UAS Dcr 2* / +, *UAS-Drs RNAi* /+. Mean and raw data points are shown for B, C, D & F. Repeated measures ANOVA for (A) p<0.0001, and (E) p<0.0001. (C) Student's t-test (D), Mann-Whitney U test (B & F), Modified Bonferroni correction following one way ANOVA for time (B) and genotype (F) (n=28-32 flies/genotype). **p <0.01, ***p <0.0001. Iso = *iso* $w^{1118}$, GrpC = Group C

### Supplementary Figure 6

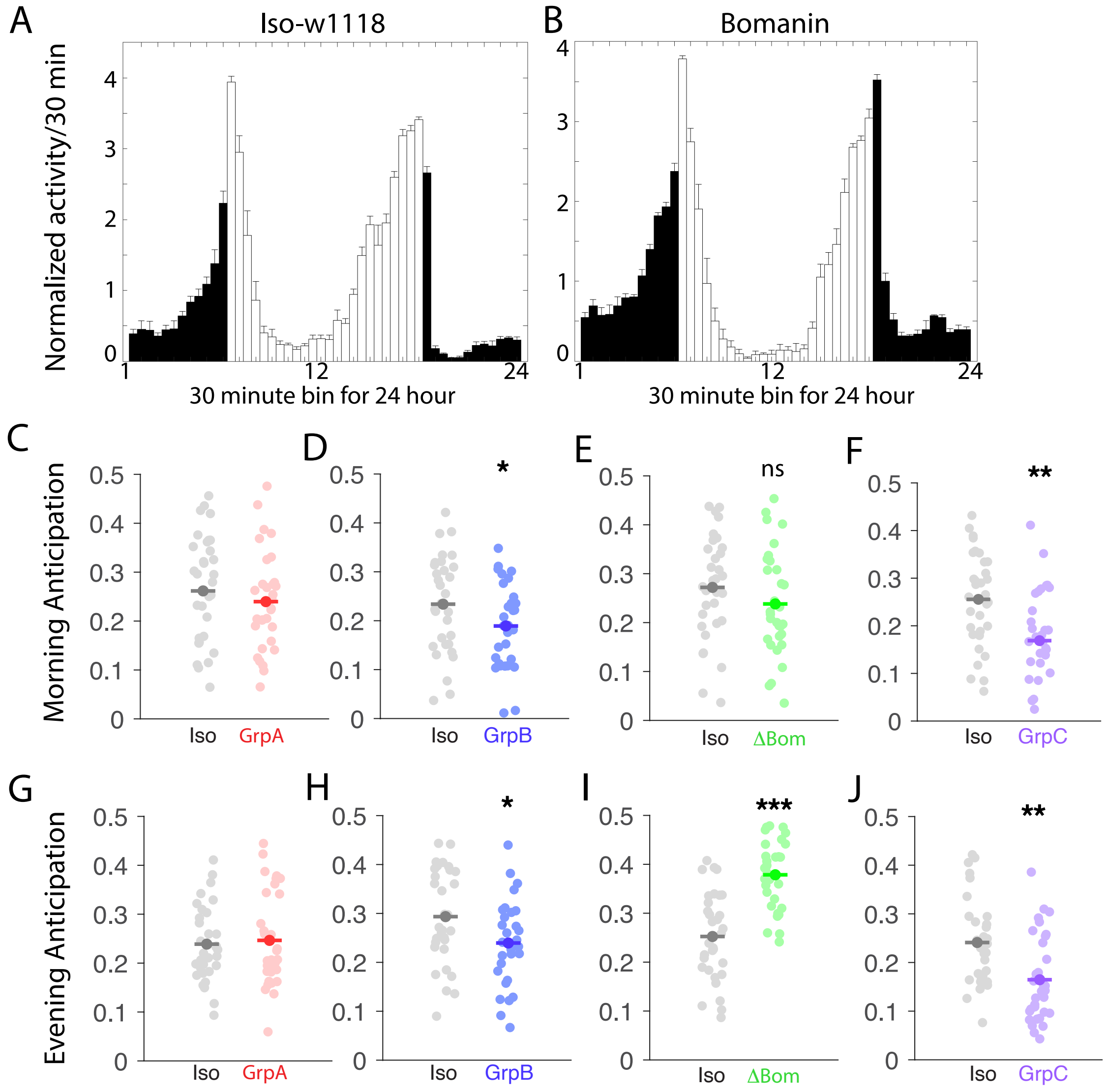


**Supplementary Fig6- Group B and Bomanin mutants are perturbed in morning and evening anticipation.** (A & B) Normalized activity/30 min plotted for morning and evening anticipation for *iso w*[1118] controls and Bomanin mutants. Mean and raw data points for morning anticipation (C- F) and evening anticipation (G- J) are shown for controls-*iso w*[1118] and respective AMP mutants. (C- J) Student's t-test comparing *iso w*[1118] controls and AMP mutants (n=24-32 flies/genotype). *p<0.05, ***p <0.0001. Iso = *iso w*[1118], GrpA = Group A, GrpB = Group B, ΔBom = Bomanins, GrpC = Group C

#### Supplementary Figure 7

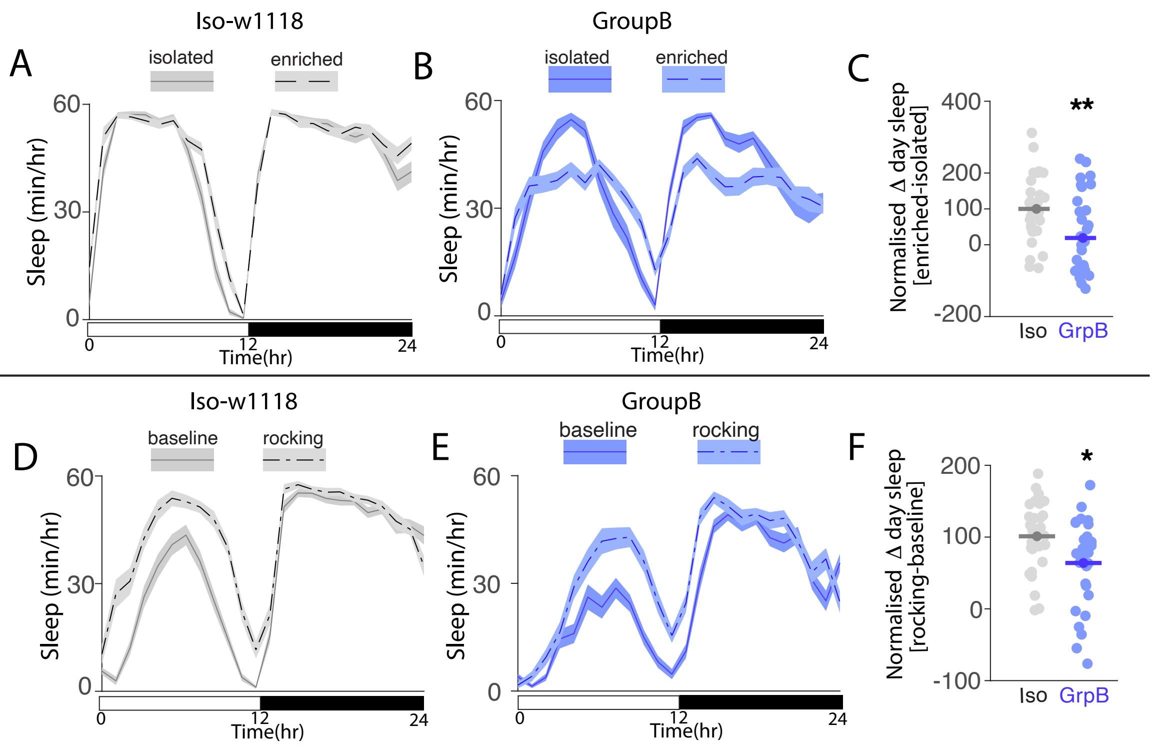


**Supplementary Fig7- Characteristics of socialization and rocking induced sleep in Group B mutants.** (A & B) Representative experiment showing sleep (mean ± SEM sleep) per hour for isolated and socially enriched *iso* $w^{1118}$ controls (A) and Group B mutants (B). Sleep is shown for 24h, with hours 0-12 as the light period, 12-24 as the dark period. Repeated measures ANOVA detected a significant effect of condition (socialization) in *iso* $w^{1118}$ controls, p= 0.00071, but not in Group B mutants, p= 0.0788. (C) Percent change in day sleep upon social enrichment for Group B normalized to *iso* $w^{1118}$ controls; mean and raw data points are shown. (D & E) Representative experiment showing sleep (mean ± SEM sleep) per hour for *iso* $w^{1118}$ controls (D) and Group B mutants (E) on baseline and after rocking. Sleep is shown for 24h, with hour 0-12 as the light period, 12-24 as the dark period. Repeated measures ANOVA detected significant effect of condition (rocking) for *iso* $w^{1118}$ controls, p= 4.06 x10-8, and Group B, p= 5.40 x10-5. (F) Percent change in day sleep upon rocking for Group B normalized to *iso* $w^{1118}$ controls, mean and raw data is shown. (C & F) Student's t-test for comparing between *iso* $w^{1118}$ controls and Group B mutants (n=22-32 flies / genotype). *p<0.05, **p <0.01.

#### Supplementary Figure 8

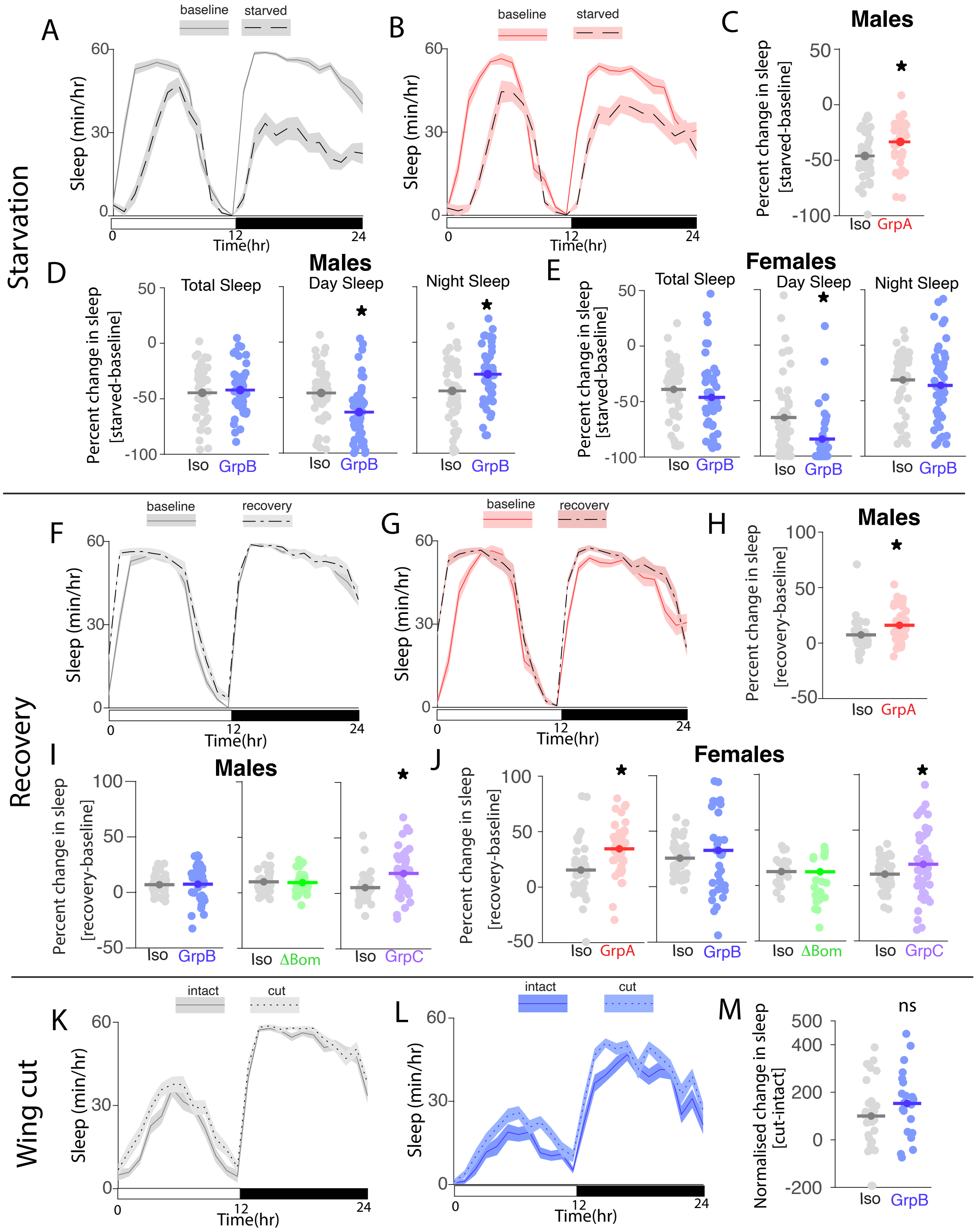

**Supplementary Fig8- Characteristics of starvation and wing-cut induced sleep-plasticity in AMP mutants.** (A & B) Representative experiment showing sleep (mean ± SEM sleep) per hour on baseline and starvation day for *iso* $w^{1118}$ controls (A) and Group A mutants (B). Sleep is shown for 24h, with hour 0-12 as the light period, 12-24 as the dark period. Repeated measures ANOVA detected significant effect of condition (starvation) for *iso* $w^{1118}$ controls, p= 9.99 x10-23, and Group A mutants, p= 3.76 x10-13. (C) Normalized change in sleep upon starvation for *iso* $w^{1118}$ controls and Group A males, mean and raw data points are shown. (D & E) Normalized change in total sleep, day sleep and night sleep upon starvation for Isow1118 controls and Group B males (D) and females (E) respectively. Mean and raw data points are shown. (F & G): Representative experiment showing sleep (mean ± SEM sleep) per hour on baseline and recovery day for *iso* $w^{1118}$ controls (F) and Group A mutants (G). Sleep is shown for 24h, with hour 0-12 as the light period, 12-24 as the dark period. Repeated measures ANOVA detected significant effect of condition (recovery following starvation) for *iso* $w^{1118}$ controls, p= 0.00027 (F), and Group A mutants, p= 1.39 x10-6 (G). (H) Percent change in sleep during recovery for *iso* $w^{1118}$ controls and Group A males. Mean and raw data points are shown. (I & J) Percent change in sleep during recovery for *iso* $w^{1118}$ controls and Group B, Bomanin and Group C males (I) and females (J) respectively. Mean and raw data points are shown.(K & L): (K) Representative experiment showing sleep (mean ± SEM sleep) per hour of flies with intact and cut wings for *iso* $w^{1118}$ controls (K) and Group B mutants (L). Sleep is shown for 24h, with hour 0-12 as the light period, 12-24 as the dark period. Repeated measures ANOVA detected significant effect of condition (wingcut) for *iso* $w^{1118}$ controls, p= 0.0116, and Group B mutants, p= 0.0022. (M) Normalized change in sleep following wing-cut for *iso* $w^{1118}$ controls and Group B females, mean and raw data points are shown. *p<0.05. Iso = *iso* $w^{1118}$, GrpA = Group A, GrpB = Group B, ∆Bom = Bomanins, GrpC = Group C

### Supplementary Figure 9

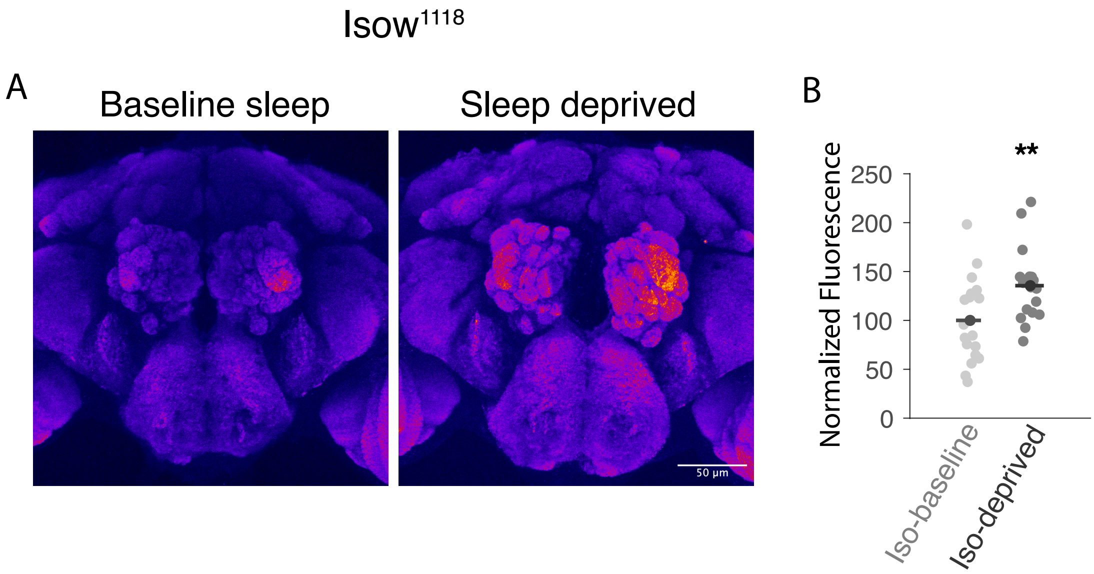


**Supplementary Fig9- Elevated BRP expression following sleep deprivation.** (A) Brains of *iso* $w^{1118}$ controls under baseline (left) and sleep deprived (right) conditions stained with mAb nc82. Z-projections from whole mount immunostaining processed using the Fire LUT in FiJi. Scale bar = 50µm. (B) Quantification of BRP expression in Isow1118 brains under baseline and sleep deprived condition. (B) Student's t test (n=20-23 brains /condition). **p <0.01.

#### Supplementary Figure 10

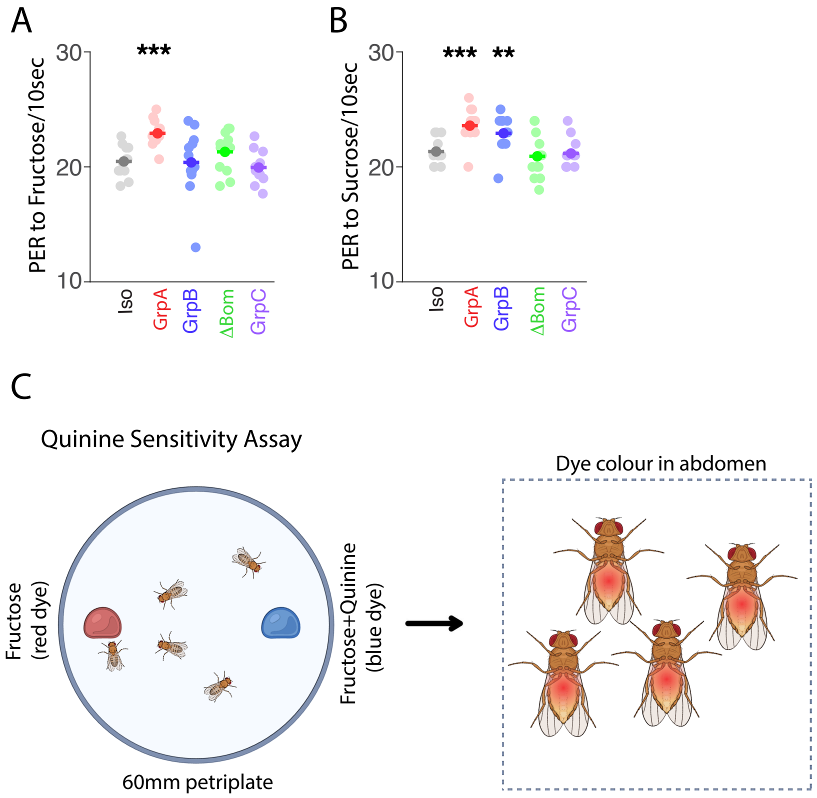


D

Table 1: Quinine senstivity of AMP mutants- feeding choice assay

| S. no | Genotype | Choice A: Fructose (50mM) | Choice B: Quinine (50mM) + Fructose (50mM) | Total flies tested | % flies avoid quinine |
|---|---|---|---|---|---|
| 1 | Iso-w1118 | 21 | 0 | 21 | 100 |
| 2 | GroupA | 27 | 0 | 27 | 100 |
| 3 | GroupB | 28 | 0 | 28 | 100 |
| 4 | ΔBomanins | 25 | 0 | 25 | 100 |
| 5 | GroupC | 28 | 0 | 28 | 100 |
| 6 | ΔBomanins (Vehicle fed) | 20 | 0 | 20 | 100 |
| 7 | ΔBomanins (Gaboxadol fed) | 26 | 0 | 26 | 100 |
| 8 | ΔBomanins (L12-D12) | 22 | 0 | 22 | 100 |
| 9 | ΔBomanins (L14-D10) | 30 | 0 | 30 | 100 |

**Supplementary Fig10- AMP mutants are not impaired in sweet or bitter taste sensation.** (A) Number of proboscis extensions to fructose presented on tarsi (100mM for 10s) to *iso* $w^{1118}$ controls and AMP mutants. Mean and raw data points are shown. (B) Number of proboscis extensions to sucrose presented on tarsi (500mM for 10s) to *iso* $w^{1118}$ controls and AMP mutants. Mean and raw data points are shown. (C) Schematic of the quinine sensitivity assay. (D) Table showing the number and percent of flies that chose fructose vs fructose+ quinine. (A & B) Modified Bonferroni correction following one way ANOVA for genotype (n=12 flies / genotype). **p <0.01, ***p <0.0001. Iso = *iso* $w^{1118}$, GrpA= Group A, GrpB= Group B, ΔBom= Bomanins, GrpC= Group C.

#### Supplementary Figure 11

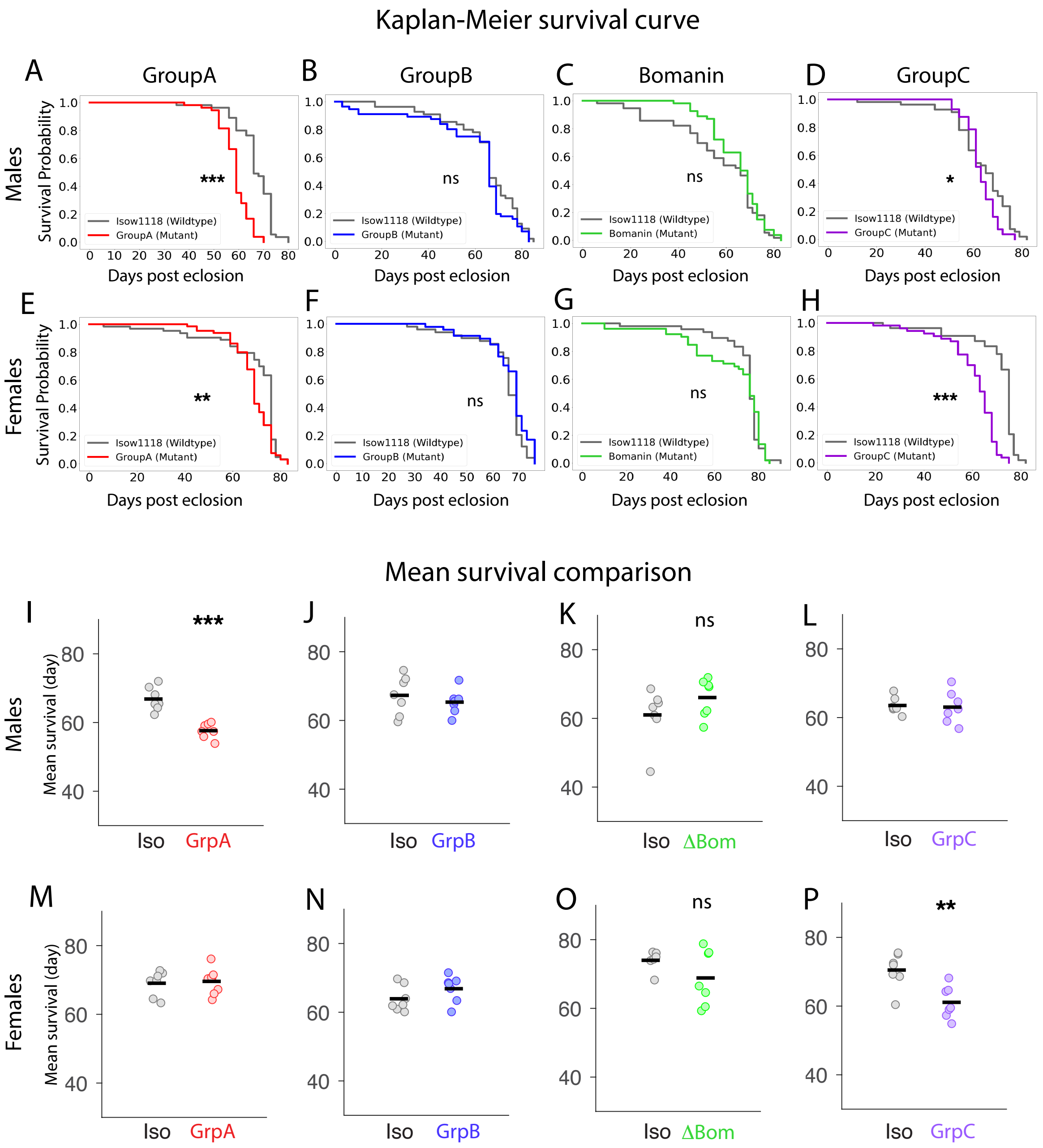


**Supplementary Fig11- Group A and Group C mutant flies have a shortened lifespan.** (A-D) Kaplan-Meier survival curves for *iso* $w^{1118}$ controls and Group A (A), Group B (B), Bomanin (C), and Group C (D) mutant male flies. (E-H) Kaplan-Meier survival curves for *iso* $w^{1118}$ controls and Group A (A), Group B (B), Bomanin (C), and Group C (D) mutant female flies. (I-L) Mean survival of 7-9 groups of *iso* $w^{1118}$ and AMP mutant males, with 7-8 flies each. (M-P) Mean survival of 7-9 groups of *iso* $w^{1118}$ and AMP mutant females, with 7-8 flies each. Mean survival of mutant group was compared to *iso* $w^{1118}$ controls. (A-H) Log-rank test, (n=50-69 flies/ genotype). (I-P) Student's t-test, (n=50-69 flies/ genotype). * $p<0.05$, ** $p<0.01$, *** $p<0.0001$. Iso = *iso* $w^{1118}$, GrpA= Group A, GrpB= Group B, ΔBom= Bomanins, GrpC= Group C

Supplementary Figure 12

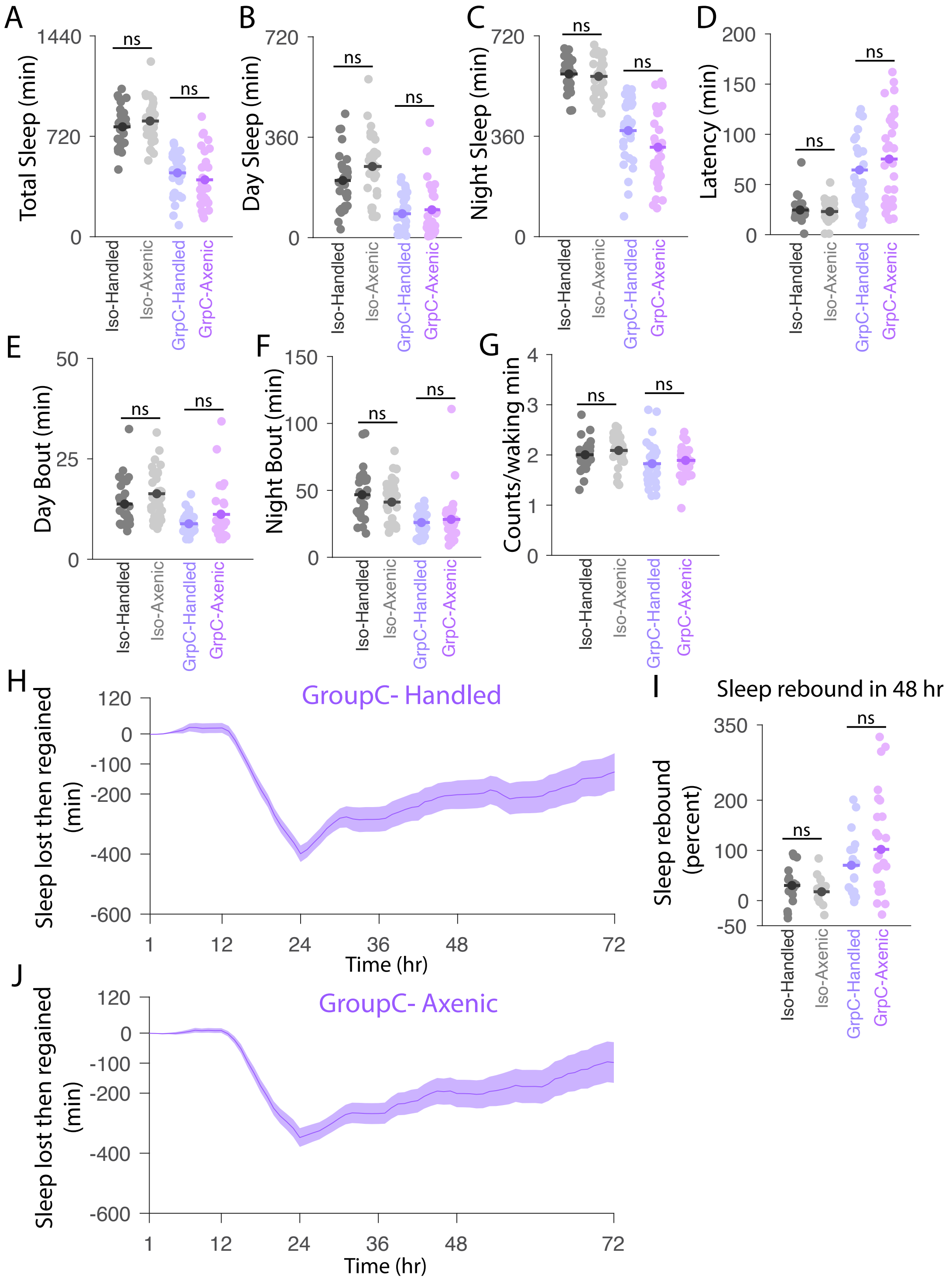


**Supplementary Fig12- Loss of function of Group C genes affects baseline sleep and sleep homeostasis independent of gut microbiota.** (A-G) Mean and raw data points are shown for total sleep (A), day sleep (B), night sleep (C), latency (D), day bout length (E), night bout length (F), and waking activity (G) of handled and axenic groups of *iso* $w^{1118}$ controls and Group C mutants. (H, J) Plot of mean ± SEM sleep lost during overnight sleep deprivation and gained during 48h of recovery for Group C-handled (H) and Group C-axenic (J) flies are shown. (I) Mean and raw data points are shown for percent sleep regained over 48 hours of recovery for handled and axenic *iso* $w^{1118}$controls and Group C mutants. Axenic group was compared to handled control for each genotype. (A, C, E, F, I) Student's t-test, (B, D, G) Mann-Whitney test, (28-32 flies/ genotype), ns-not significant. Iso = *iso* $w^{1118}$ , GrpC= Group C

### Supplementary Figure 13

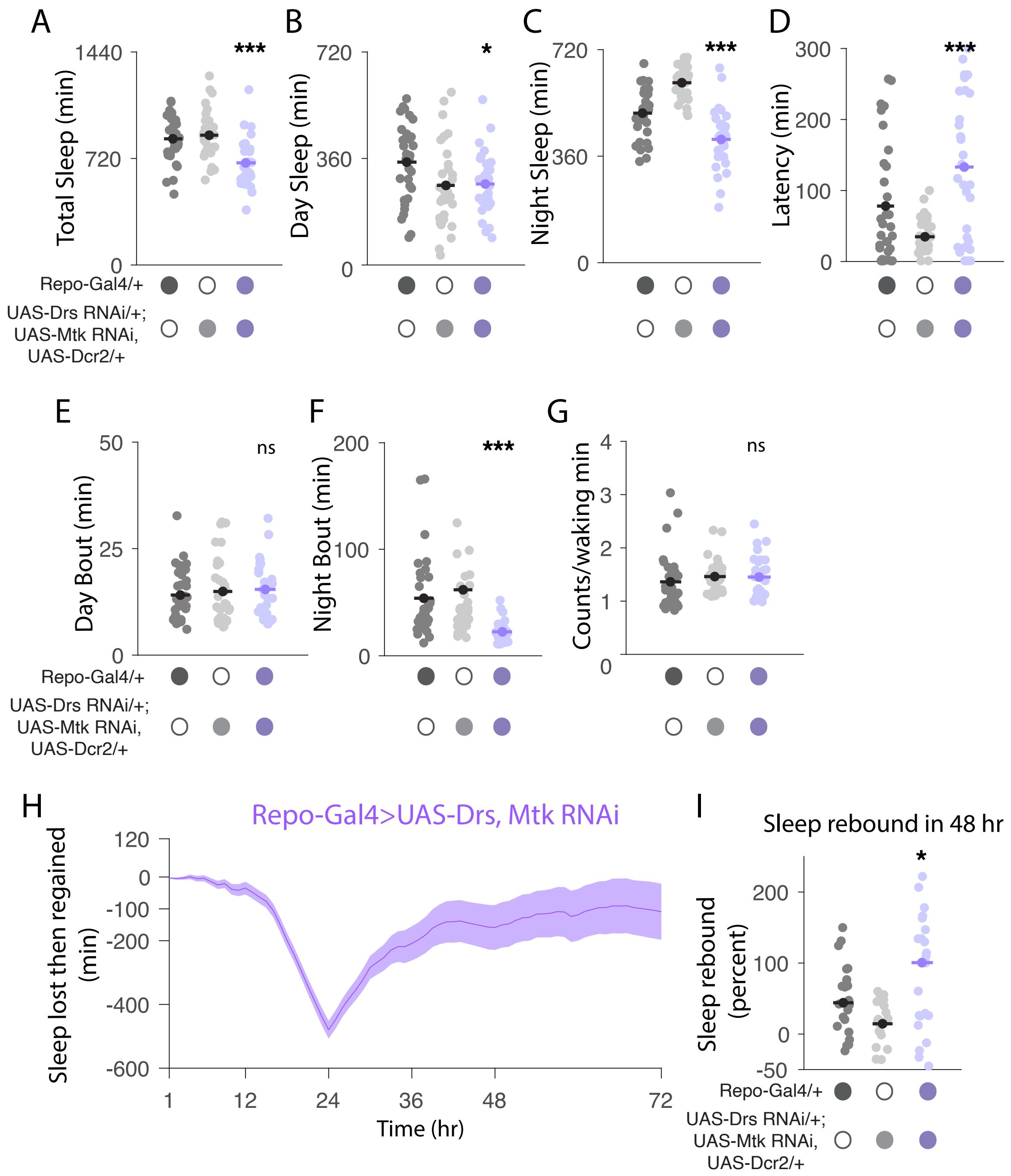


**Supplementary Fig13- Glial knockdown of Metchnikowin and Drosomycin mimics the reduced baseline sleep and exaggerated sleep rebound phenotypes of Group C mutants.** (A-G) Mean and raw data points are shown for total sleep (A), day sleep (B), night sleep (C), latency (D), day bout length (E), night bout length (F), and waking activity (G) for *Repo-Gal4> UAS-Drs RNAi* ; *UAS-Mtk RNAi UAS Dcr2* and their genetic controls *Repo-Gal4*/+ and *UAS-Drs RNAi*/+; *UAS-Mtk RNAi UAS Dcr2* / +;. (H) Plot of mean ± SEM sleep lost during overnight sleep deprivation and gained during 48h of recovery for *Repo-GAL4> UAS-Drs RNAi* ; *UAS-Mtk RNAi UAS Dcr2*. (I) Mean and raw data points are shown for percent sleep regained over 48 hours of recovery for *Repo GAL4> UAS-Drs RNAi* ; *UAS-Mtk RNAi UAS Dcr2* and the genetic controls, *Repo GAL4*/+ and *UAS-Drs RNAi*/+; *UAS-Mtk RNAi UAS Dcr2* / +. (A, B, C) Modified Bonferroni correction following one way ANOVA for genotype. (D-G, I) Dunn's multiple comparisons following Kruskal-Wallis ANOVA (n=30-32 flies/ genotype), * p<0.05, ** p <0.01, *** p <0.0001.

#### Supplementary Figure 14

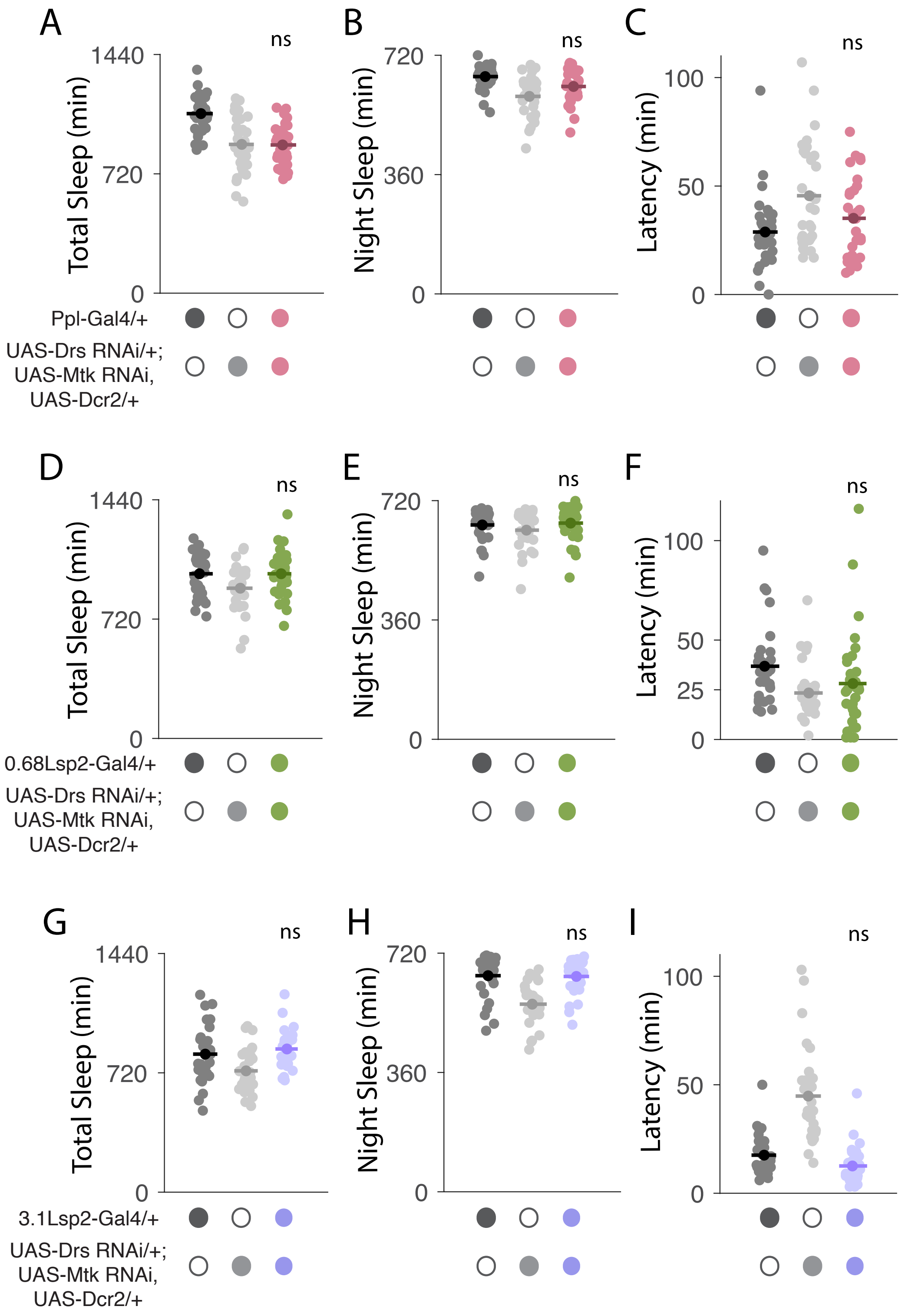


**Supplementary Fig14- Fat-body knockdown of Metchnikowin and Drosomycin does not perturb baseline sleep.** (A-C) Mean and raw data points are shown for total sleep (A), night sleep (B), and latency (C) for *Ppl GAL4> UAS Drs RNAi* ; *UAS Mtk RNAi UAS Dcr2* and their genetic controls *Ppl GAL4/+* and *UAS Drs RNAi/+*; *UAS Mtk RNAi, UAS Dcr2* / +. (D-F) Mean and raw data points are shown for total sleep (D), night sleep (E), and latency (F) for *0.68Lsp2 GAL4> UAS Drs RNAi* ; *UAS Mtk RNAi UAS Dcr2* i and their genetic controls *0.68Lsp2 GAL4/+* and *UAS Drs RNAi/+*; *UAS Mtk RNAi UAS Dcr2* / +. (G-I) Mean and raw data points are shown for total sleep (A), night sleep (B), and latency (C) for *3.1Lsp2-GAL4> UAS Drs RNAi* ; *UAS Mtk RNAi UAS Dcr2* and their genetic controls *3.1Lsp2 GAL4/+* and *UAS Drs RNAi/+*; *UAS Mtk RNAi UAS Dcr2* / +. (A, D, G) Modified Bonferroni correction following one way ANOVA for genotype. (B, C, E, F, H, I) Dunn's multiple comparisons following Kruskal-Wallis ANOVA (n=28-32 flies/ genotype), ns- not significant.

### Supplementary Figure 15

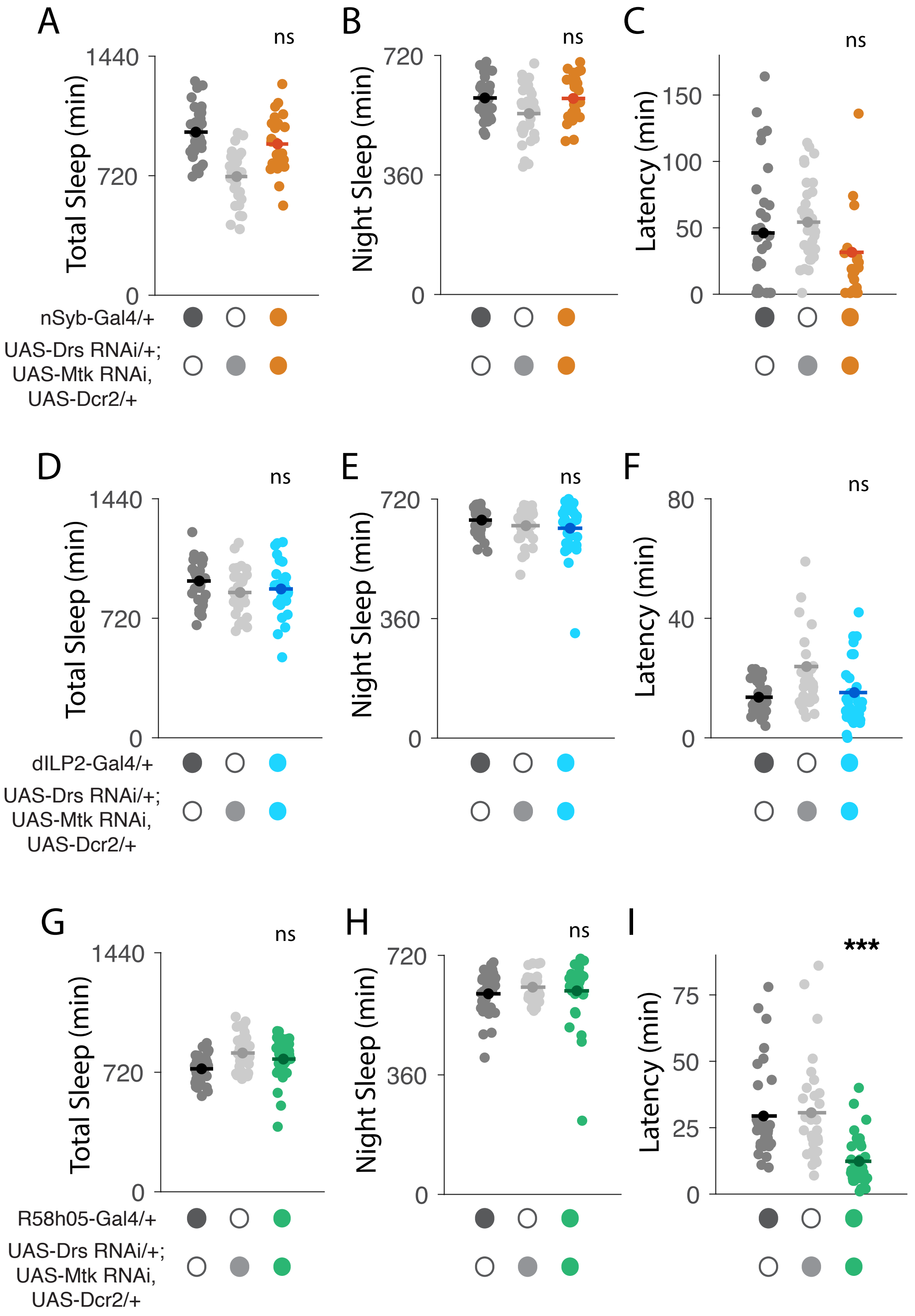


**Supplementary Fig15- Pan-neuronal and subset-specific knockdown of Metchnikowin and Drosomycin does not mimic Group C sleep phenotypes.** (A-C) Mean and raw data points are shown for total sleep (A), night sleep (B), and latency (C) for *nSyb GAL4> UAS Drs RNAi* ; *UAS-Mtk RNAi UAS Dcr2* and their genetic controls *nSyb GAL4/+* and *UAS Drs RNAi/+*; *UAS Mtk RNAi UAS Dcr2* / +. (D-F) Mean and raw data points are shown for total sleep (D), night sleep (E), and latency (F) for *dILP2 GAL4> UAS Drs RNAi* ; *UAS Mtk RNAi UAS Dcr2* and their genetic controls *dILP2 GAL4/+* and *UAS Drs RNAi/+*; *UAS Mtk RNAi UAS Dcr2* / +. (G-I) Mean and raw data points are shown for total sleep (A), night sleep (B), and latency (C) for *R58H05 GAL4> UA -Drs RNAi* ; *UAS Mtk RNAi UAS Dcr2* and their genetic controls *R58H05 GAL4/+ UAS Drs RNAi/+*; *UAS-Mtk RNAi UAS Dcr2* / +. (A, B, D) Modified Bonferroni correction following one way ANOVA for genotype. (C, E, F, G, H, I) Dunn's multiple comparisons following Kruskal-Wallis ANOVA (n=26-32 flies/ genotype), ns- not significant, *** p <0.0001.

#### Supplementary Figure 16

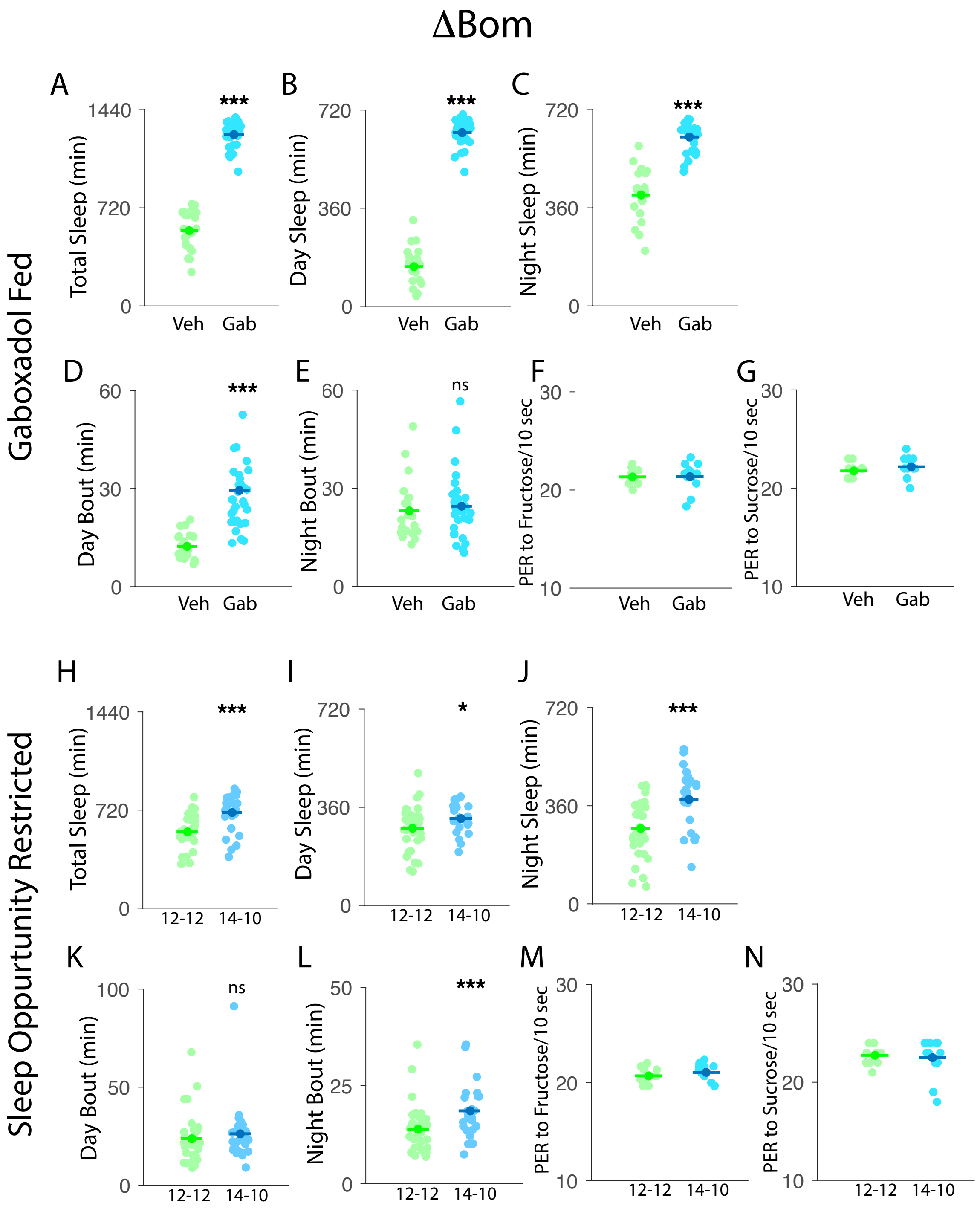


**Supplementary Fig16- Sleep characteristics and sugar sensing in gaboxadol fed and sleep opportunity restricted Bomanin mutants.** (A-G) Mean and raw data points are shown for total sleep (A), day sleep (B), night sleep (C), day bout length (D), night bout length (E), number of proboscis extensions to fructose (100mM for 10s) presented on tarsi, and (F) number of proboscis extensions to sucrose (500mM for 10s) presented on tarsi (G) for vehicle fed (veh) and gaboxadol fed (gab) Bomanin mutant flies. Sleep characteristics of gaboxadol fed mutants were compared to vehicle fed control mutants. (H-N) Mean and raw data points are shown for total sleep (H), day sleep (I), night sleep (J), day bout length (K), night bout length (L), number of proboscis extensions to fructose (100mM for 10s) presented on tarsi, and (M) number of proboscis extensions to sucrose (500mM for 10s) presented on tarsi (N) for Bomanin mutant flies maintained on a standard 12:12 Light: Dark schedule and sleep opportunity restricted (L14-D10) Bomanin mutant flies. Sleep characteristics of sleep restricted mutants were compared to flies maintained on a standard 12:12 LD cycle. (A-N) Student's t-test (n=24-32 flies/condition). *p<0.05, ***p <0.0001.
